# Animal, fungi, and plant genome sequences harbour different non-canonical splice sites

**DOI:** 10.1101/616565

**Authors:** Katharina Frey, Boas Pucker

## Abstract

Most protein encoding genes in eukaryotes contain introns which are interwoven with exons. After transcription, introns need to be removed in order to generate the final mRNA which can be translated into an amino acid sequence. Precise excision of introns by the spliceosome requires conserved dinucleotides which mark the splice sites. However, there are variations of the highly conserved combination of GT at the 5’ end and AG at the 3’ end of an intron in the genome. GC-AG and AT-AC are two major non-canonical splice site combinations which have been known for years. During the last years, various minor non-canonical splice site combinations were detected with numerous dinucleotide permutations. Here we expand systematic investigations of non-canonical splice site combinations in plants to all eukaryotes by analysing fungal and animal genome sequences. Comparisons of splice site combinations between these three kingdoms revealed several differences such as a substantially increased CT-AC frequency in fungal genome sequences. Canonical GT-AG splice site combinations in antisense transcripts could be one explanation for this observation. In addition, high numbers of GA-AG splice site combinations were observed in *Eurytemora affinis* and *Oikopleura dioica*. A variant in one U1 snRNA isoform might allow the recognition of GA as 5’ splice site. In depth investigation of splice site usage based on RNA-Seq read mappings indicates a generally higher flexibility of the 3’ splice site compared to the 5’ splice site across animals, fungi, and plants.

## Introduction

Splicing, the removal of introns after transcription, is an essential step during the generation of mature mRNAs in eukaryotes. This process allows variation which provides the basis for quick adaptation to changing conditions^1,2^. Alternative splicing, e.g. skipping exons, usage of alternative 5’ or 3’ splice sites and the retention of introns, results in an enormous diversity of synthesized proteins and therefore substantially expands the diversity of products encoded in eukaryotic genomes^3–6^.

The full range of functions as well as the evolutionary relevance of introns are still under discussion^7^. However, introns are energetically expensive for the cell to maintain as the transcription of introns costs time and energy and the removal of introns has to be exactly regulated^8^. Dinucleotides at both intron/exon borders mark the splice sites and are therefore highly conserved^9^. GT at the 5’ end and AG at the 3’ end of an intron form the canonical splice site combination on DNA level. More complexity arises through non-canonical splice site combinations, which deviate from the highly conserved canonical one. Besides the major non-canonical splice site combinations GC-AG and AT-AC, several minor non-canonical splice site combinations have been detected before^9,10^.

Furthermore, the position of introns in homologous genes across organisms, which diverged 500-1500 million years ago, are not conserved^11^. In addition, many intron sequences mutate at a higher rate due to having much less of an impact on the reproductive fitness of an organism compared to a mutation located within an exon^12^. These factors, along with the existence of several non-canonical splice sites, make the complete prediction of introns, even in non-complex organisms like yeast, almost impossible^13,14^. Moreover, most introns which can be predicted computationally still lack experimental support^15^.

Splice sites are recognised during the splicing process by a complex of snRNAs and proteins, the spliceosome^16^. U2-spliceosome and U12-spliceosome are two subtypes of this complex which comprise slightly different proteins with equivalent functions^17–19^. Although the terminal dinucleotides are important for the splicing process, these splice sites are not sufficient to determine which spliceosome is processing the enclosed intron^20^. This demonstrates the complexity of the splicing process which involves additional signals present in the DNA. Even though multiple mechanisms could explain the splicing process, the exact mechanism of non-canonical splicing is still not completely resolved^5^.

Branching reaction and exon ligation are the two major steps of splicing^21,22^. In the branching reaction, the 2’-hydroxyl group of the branchpoint adenosine initiates an attack on the 5’-phosphate of the donor splice site^23,24^. This process leads to the formation of a lariat structure. Next, the exons are ligated and the intron is released through activity of the 3’-hydroxyl group of the 5’ exon at the acceptor splice site^21^.

Previous in-depth analyses of non-canonical splice sites in fungi and animals were often focused on a single or a small number of species^9,25,26^. Several studies focused on canonical GT-AG splice sites but neglected non-canonical splice sites^27,28^. Our understanding of splice site combinations is more developed in plants compared to other kingdoms^10,29–33^. Previous works reported 98 % GT-AG splice site combinations in fungi^25^, 98.7 % in plants,^10^ and 98.71 % in animals^9^. Consequently, the proportion of non-canonical splice sites, other than the canonical splice site GT-AG, is around or below 2 %^9,10,25^. To the best of our knowledge, it is not known if the value reported for mammals is representative for all animals. Non-canonical splice site combinations can be divided into major non-canonical GC-AG and AT-AC combinations and the minor non-canonical splice sites which are all other dinucleotide combinations at the terminal intron positions. The combined frequency of all minor non-canonical splice site combinations is low e.g. 0.09 % in plants, but still exceeds the frequency of the major non-canonical AT-AC splice sites^10^. Despite this apparently low frequency, non-canonical splice site combinations have a substantial impact on gene products, especially on exon-rich genes^10^. Over 40 % of plant genes with exactly 40 exons are affected^10^.

Consideration of non-canonical splice sites is important for gene prediction approaches, because *ab initio* identification of these splice sites is computationally extremely expensive and therefore rarely applied^29^. Moreover, as many human pathogenic mutations occur at the donor splice site^34^, it is of great interest to understand the occurrence and usage of non-canonical splice sites. Therefore, several non-canonical splice sites containing AG at the acceptor site were investigated in human fibroblasts^34^. Alongside this, fungi are interesting due to pathogenic properties and importance in the food industry^35^. Since splicing leads to high protein diversity^3–6^, the analysis of splicing in fungi is important with respect to biotechnological applications e.g. development of new products.

Non-canonical splice sites are frequently considered as artifacts^36^ and therefore excluded from analyses^27,28^. Further, RNA editing of GT-AA to GT-AG splice sites on RNA level is possible^37^. This leads to the transformation of non-canonical splice site combinations into canonical ones. Previous studies supported minor non-canonical splice site combinations in single or few species^9,25,26^ and systematically across plants^10,29–33^. In this study, a collection of annotated genome sequences from 130 fungi and 489 animal species was screened for canonical and non-canonical splice site combinations in representative transcripts. RNA-Seq data sets were harnessed to identify biologically relevant and actually used splice sites based on the available annotation. Non-canonical splice site combinations, which appeared at substantially higher frequency in a certain kingdom or species, were analysed in detail. As knowledge about splice sites in plants was available from previous investigations^10,29^, a comparison between splice sites in fungi, animals and plants was performed.

## Results and Discussion

### Analysis of non-canonical splice sites

In total, 64,756,412 (Supplementary Data S1) and 2,302,340 (Supplementary Data S2) splice site combinations in animals and fungi, respectively, were investigated based on annotated genome sequences (Supplementary Data S3 and Supplementary Data S4). The average frequency of the canonical splice site combination GT-AG is 98.3 % in animals and 98.7 % in fungi, respectively. These values exceed the 97.9 % previously reported for plants^10^, thus indicating a generally higher frequency of non-canonical splice site combinations in plants. As previously speculated^10^, a generally less accurate splicing system in plants could be an adaptation to changing environments through the generation of a larger transcript diversity. Since most plants are not able to change their geographic location, the tolerance for unfavorable conditions should be stronger than in animals. The lower proportion of non-canonical splice site combinations in fungi compared to animals seems to contradict this hypothesis. However, the genome size and complexity needs to be taken into account here. Based on the available assemblies, the average animal genome is significantly larger than the average fungal genome (Mann-Whitney U-Test; p=5.64e-68) (Supplementary Data S5). Although the average animal genome sequence (median=998 Mbp) is longer than the plant average (median=467 Mbp), plant genome sequences harbour more non-canonical splice sites (Supplementary Data S6, Supplementary Data S7^10^).

Average percentages of the most important splice site combinations were summarized per kingdom and over all analysed genomes (Table 1). The number of canonical and non-canonical splice site combinations per species was also summarized (Supplementary Data S8 and Supplementary Data S9). A higher percentage of non-canonical splice sites was observed in animals in comparison to fungi. Several species strongly exceeded the average values for major and minor non-canonical splice sites. The fungal species *Meyerozyma guilliermondi* shows approximately 6.67 % major and 13.33 % minor non-canonical splice sites. *Eurytemora affinis* (copepod) and *Oikopleura dioica* (tunicate) reveal approximately 10 % minor non-canonical splice sites. In summary, the observed frequencies of canonical and major non-canonical splice site combinations are similar to the pattern previously reported for plants^10^, but some essential differences and exceptions were found in animals and fungi. Previous studies already revealed that non-canonical splice site combinations are not just the result of sequencing errors^9,10,29,38^. Here, we investigated the position of sequence variants in plants, fungi, and animals with respect to splice sites. The average frequency of sequence variants at splice sites is far below 1% (Supplementary Data S10). Although non-canonical splice sites are generally more likely to harbour sequence variants than canonical ones, these sequence variants can only account for a very small proportion of non-canonical splice site combinations.

**Table 1:** Splice site combination frequencies in animals, fungi, and plants. Only the most frequent combinations are displayed here and all minor non-canonical splice site combinations are summarized as one group (“others”). A full list of all splice site combinations is available (Supplementary Data S1 and Supplementary Data S2).

|  | GT-AG | GC-AG | AT-AC | others |
| --- | --- | --- | --- | --- |
| animals | 98.334 % | 0.983 % | 0.106 % | 0.577 % |
| fungi | 98.715 % | 1.009 % | 0.019 % | 0.257 % |
| plants | 97.886 % | 1.488 % | 0.092 % | 0.534 % |
| all | 98.265 % | 1.074 % | 0.101 % | 0.560 % |

Different properties of the genome sequences of all investigated species were analysed to identify potential explanations for the splice site differences (Supplementary Data S6 and Supplementary Data S7). In fungi, the average number of introns per gene is 1.49 and the average GC content is 47.1 % (±7.39; s.d.). In animals, each gene contains on average 6.95 introns and the average GC content is 39.4 % (±3.87; s.d.). The average number of introns per gene in plants is 4.15 and the average GC content 36.3 % (±8.84; s.d.). This difference in the GC content between fungi and animals/plants could be associated with the much lower frequency of AT-AC splice site combinations and the higher frequency of CT-AC splice site combinations in fungi (Figure 1). CT-AC has a higher GC content than the AT rich AT-AC splice site combination. A generally higher GC content could result in the higher GC content within splice site combinations due to the overall mutation rates in these species.

**Figure 1:**
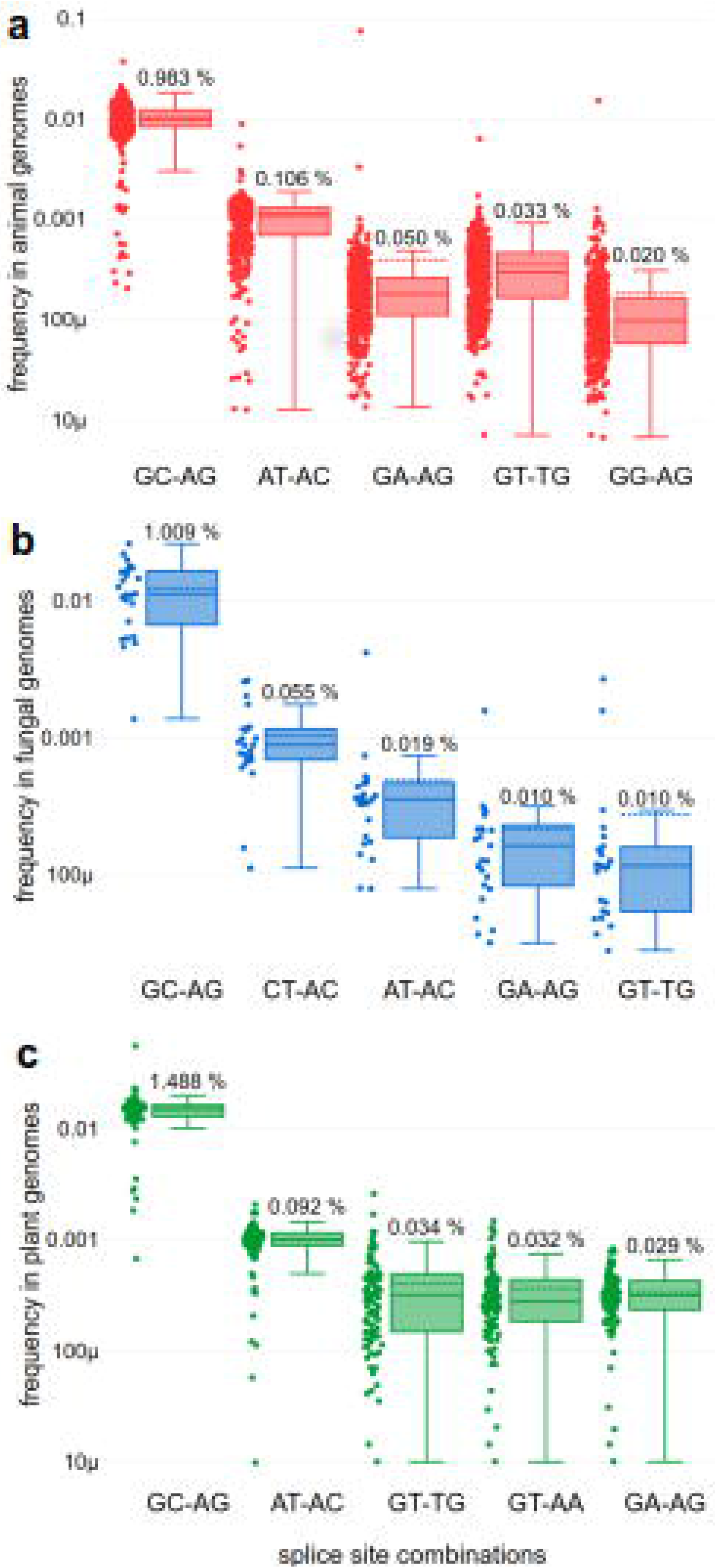
Frequencies of non-canonical splice site combinations in animals, fungi, and plants. The frequency of non-canonical splice site combinations across the 489 animal (red), 130 fungal (blue) and 121 plant (green) genomes is shown. Normalization of the absolute number of each splice site combination was performed per species based on the total number of splice sites. The frequency of the respective splice site combination of each species is shown on the left-hand side and the percentage of the respective splice site combination on top of each box plot. The dashed line represents the mean frequency of the respective splice site combination over all investigated species. The box plots are ordered (from left to right) according to the mean frequency.

A comparison of the genome-wide GC content to the GC content of all splice sites revealed a weak correlation in the analysed fungi (r≈0.236, p≈0.008). Species with a high genomic GC content tend to show a high GC content in the splice site combinations in the respective species. A similar correlation was found in plant (r≈0.403, p≈4.505e-06) and animal species (r≈0.434, p≈7.866e-24) (Supplementary Data S11). Additionally, the GC content in fungal genomes is substantially exceeding the average GC content of plant and animal genomes. Since genomic GC content and intronic GC content strongly correlate (animals: r≈0.968, p≈2.357e-292, plants: r≈0.974, p≈8.987e-79 and fungi: r≈0.950, p≈2.800e-66), the results obtained in the analysis above are representative for both methods of GC content calculation (Supplementary Data S11). Since splicing of U12 introns, which often show the major non-canonical AT-AC splice site combination, requires the presence of the minor U12 spliceosome, we screened the genome sequences of all investigated species for components of this spliceosome. As differences in the genome sequence completeness and continuity as well as sequence divergent from bait sequences can impact the results, we only state the presence of the U12 spliceosome in some species while the absence in the remaining species cannot be demonstrated. The comparison of annotated AT-AC splice site combinations between species with and without the minor U12 spliceosome revealed significantly higher numbers of this major non-canonical splice site combination in species with U12 spliceosomes (Mann-Whitney U-Test: p≈3.8e-12 (plants) and p≈1.8e-15 (animals)). Although many fungi are known to have a minor U12 spliceosome^19^ we only detected corresponding RNA genes in one species (*Cutaneotrichosporon oleaginosum*) and thus refrained from any conclusions about the situation in fungi.

The most frequent non-canonical splice site combinations show differences between animals, fungi, and plants (Figure 1). In fungal genome sequences, the splice site CT-AC is substantially more frequent than the splice site combination AT-AC. Regarding the splice site combination GA-AG in animal genome sequences, two outliers are clearly visible: *E. affinis* and *O. dioica* show more GA-AG splice site combinations than GC-AG splice site combinations.

Despite overall similarity in the pattern of non-canonical splice site combinations between kingdoms, specific minor non-canonical splice site combinations were identified at much higher frequency in some fungal and animal species. First, RNA-Seq data was harnessed to validate these unexpected splice site combinations. Next, the frequencies of selected splice site combinations across all species of the respective kingdom were calculated. The correlation between the size of the incorporated RNA-Seq data sets and the number of supported splice sites was examined as well (Supplementary Data S12). In animals, there is a correlation (r≈0.417, p≈0.022) between number of supported splice sites and total number of sequenced nucleotides in RNA-Seq data. For fungi, no correlation between number of supported splice sites and size of the RNA-Seq data sets could be observed. It is important to note that the number of available RNA-Seq data sets from fungi was substantially lower.

Further, analysis of introns with canonical and non-canonical splice site combinations, respectively, revealed that a higher number of introns is associated with a higher proportion of non-canonical splice sites (Supplementary Data S13).

### High diversity of non-canonical splice sites in animals

Kupfer *et al*. suggested that splicing may differ between fungi and vertebrates^25^. Our results indicate substantial differences in the diversity of splice site combinations other than GT-AG and GC-AG in fungi (H’≈0.0277) and animals (H’≈0.0637) (Kruskal-Wallis: p≈0). Besides the overall high proportion of minor non-canonical splice sites (Table 1), differences between species are high (Figure 1). The slightly higher interquartile range of splice site combination frequencies in animal species and especially in plant species (Figure 1A and C), together with the relatively high frequency of “other” splice sites in animals and plants (Table 1) suggest more variation of splice sites in the kingdoms of animals and plants compared to the investigated fungal species. Thus, the high diversity of splice sites could be associated with the higher complexity of animal and plant genomes. In addition, the difference in prevalence between the major non-canonical splice site combination GC-AG and minor non-canonical splice site combinations is smaller in animals compared to fungi and plants (Figure 1).

GA-AG is a frequent non-canonical splice site combination in some animal species. Two species, namely *E. affinis* and *O. dioica*, showed a much higher abundance of GA-AG splice site combinations compared to the other investigated species (Figure 1A). RNA-Seq reads support 5,795 (22,866 % (average)) of all GA-AG splice site combinations of both species. GA-AG splice sites are supported in all analysed species with a slightly lower frequency of 19.032 %. In *E. affinis* and *O. dioica*, the number of the GA-AG splice site combination exceeds the number of the major non-canonical splice site combination GC-AG.

For *E. affinis*, the high frequency of the GA-AG splice site combinations was described previously when GA-AG was detected in 36 introns^39^. We quantified the proportion of GA-AG splice site combinations to 3.2 % (5,345) of all 166,392 supported splice site combinations in this species. The donor splice site GA is flanked by highly conserved upstream AG and a downstream A (Figure 2). Both species, *E. affinis* (Figure 2A,B) and *O. dioica* (Figure 2C,D), show striking similarities at several positions of donor and acceptor splice sites in addition to the GA-AG dinucleotides. As the arthropod *E. affinis*^40^ and the chordate *O. dioica*^41^ belong to different phyla, the conservation of sequences flanking the donor and acceptor splice sites and the ability to splice GA-AG introns might be explained by convergent evolution or rather an ancestral trait which was only kept in a few species, including *E. affinis* and *O. dioica*.

**Figure 2:**
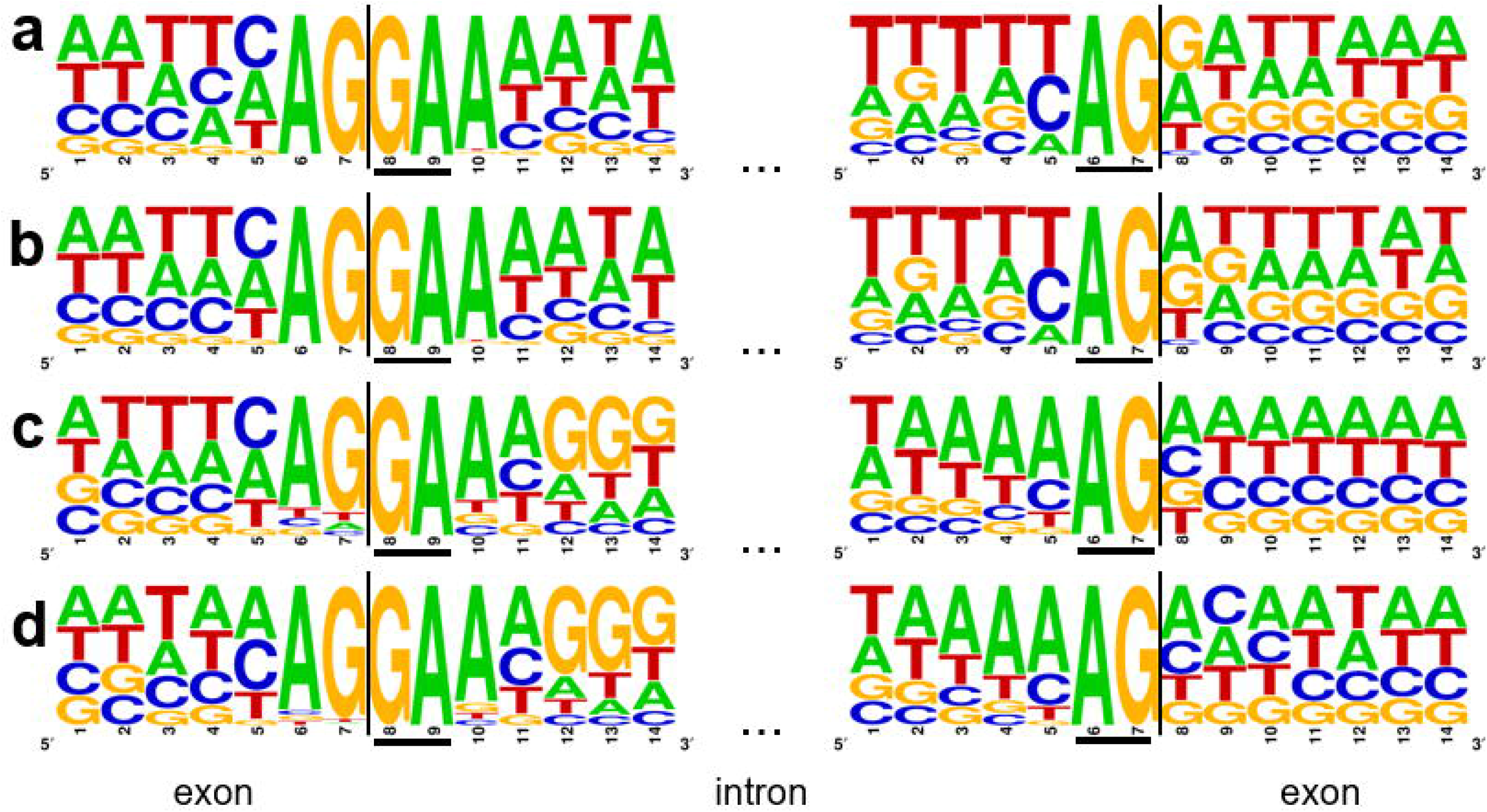
Flanking positions of GA-AG splice site combinations in *Eurytemora affinis* (a,b) and *Oikopleura dioica* (c,d). All splice site combinations (a,c) as well as all 5,795 with RNA-Seq data supported splice site combinations (b,d) of these two species were investigated. Seven exonic and seven intronic positions are displayed at the donor and acceptor splice sites. Underlined bases represent the terminal dinucleotides of the intron i.e. the donor and acceptor splice site.

Possible mechanisms which could explain these GA-AG splice site combinations are RNA editing or template switching by a reverse transcriptase. A high GC content of non-canonical splice site combinations, which is not valid for GA-AG splice sites, could facilitate the formation of secondary structures ultimately leading to template switching^36^. However, RNA editing can lead to the formation of canonical splice sites on RNA level even though a non-canonical splice site is present on DNA level^37^.

Efficient splicing of the splice site combination GA-AG was detected in human fibroblast growth factor receptor genes^42^. Further, it was suggested that this splicing event is, among other sequence properties, dependent on a canonical splice site six nucleotides upstream^42^, which does not exist in the genome sequence of the species investigated here (Figure 2). An analysis of all five potential U1 snRNAs in *E. affinis* did reveal one single nucleotide polymorphism in the binding site of the 5’ splice site from C to U at position eight in one of these U1 snRNAs. This could result in the binding of pre-mRNAs originating from A/GATAAGT instead of AG/GTAAGT (Figure 3)^39,43^. Although this would imply an elegant way for the splicing of GA-AG splice sites, the same variation was also detected in putative human U1 snRNAs. Therefore, another mechanism or additional factor might be causal for splicing of introns containing the GA-AG splice site combination. A modified copy of spliceosomal components is a likely explanation for the observed GA-AG splice site combination. However, no higher amplification ratio of spliceosome parts was observed in *E. affinis* and *O. dioica* compared to other animal species.

**Figure 3:**
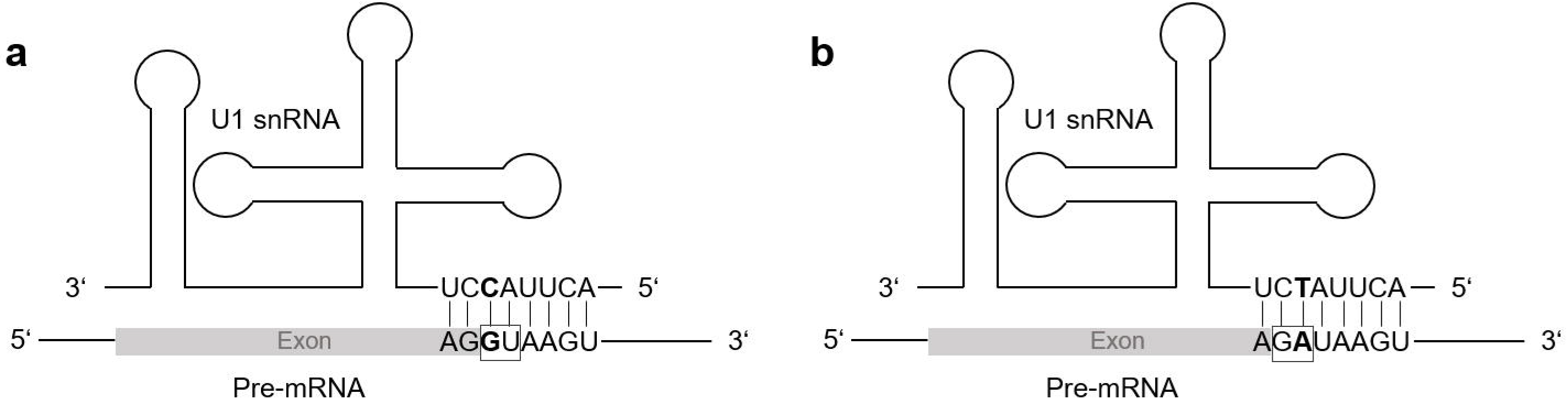
Hypothetical binding of the U1 snRNA to the pre-mRNA. A) Binding sequence of the canonical U1 snRNA to the canonical 5’ splice site GU (GT on DNA). B) Hypothetical binding sequence of the non-canonical U1 snRNA (C>T) to the non-canonical 5’ splice site GA.

### CT-AC is a frequent splice site combination in fungal annotations

Although the general frequency pattern of fungal splice site combinations is similar to plants and animals, several fungal species displayed a high frequency of minor non-canonical CT-AC splice site combinations. This co-occurs with a lower frequency of AT-AC splice site combinations. Non-canonical splice sites in fungi were, so far, only described in studies which focussed on a single or a few species. An analysis in the fungus-like microorganism *Phytophthora sojae*^44,45^, revealed 3.4 % non-canonical splice site combinations GC-AG and CT-AC^46^. Our findings indicate, that the minor non-canonical splice site combination CT-AC occurs with a significantly (Mann-Whitney U-Test; p≈0.00035) higher frequency in the annotation of fungal genome sequences than the major non-canonical splice site combination AT-AC. In contrast, the frequency of AT-AC in animals (p≈9.560e-10) and plants (p≈5.464e-24) exceeds the CT-AC frequency significantly (Figure 4A). For the splice site combination CT-AC a sequence logo, which shows the conservation of this splice site in four selected species, was designed (Figure 4B). In summary, we conclude that CT-AC is a major non-canonical splice site combination in fungi, while AT-AC is not. The highest frequencies of the splice site combination CT-AC, supported by RNA-Seq reads, were observed in *Alternaria alternata, Aspergillus brasiliensis, Fomitopsis pinicola*, and *Zymoseptoria tritici* (approx. 0.08 - 0.09 %). As AT-AC was described as major non-canonical splice site, these findings might indicate a different splice site pattern in fungi compared to animals and plants (Figure 4).

**Figure 4:**
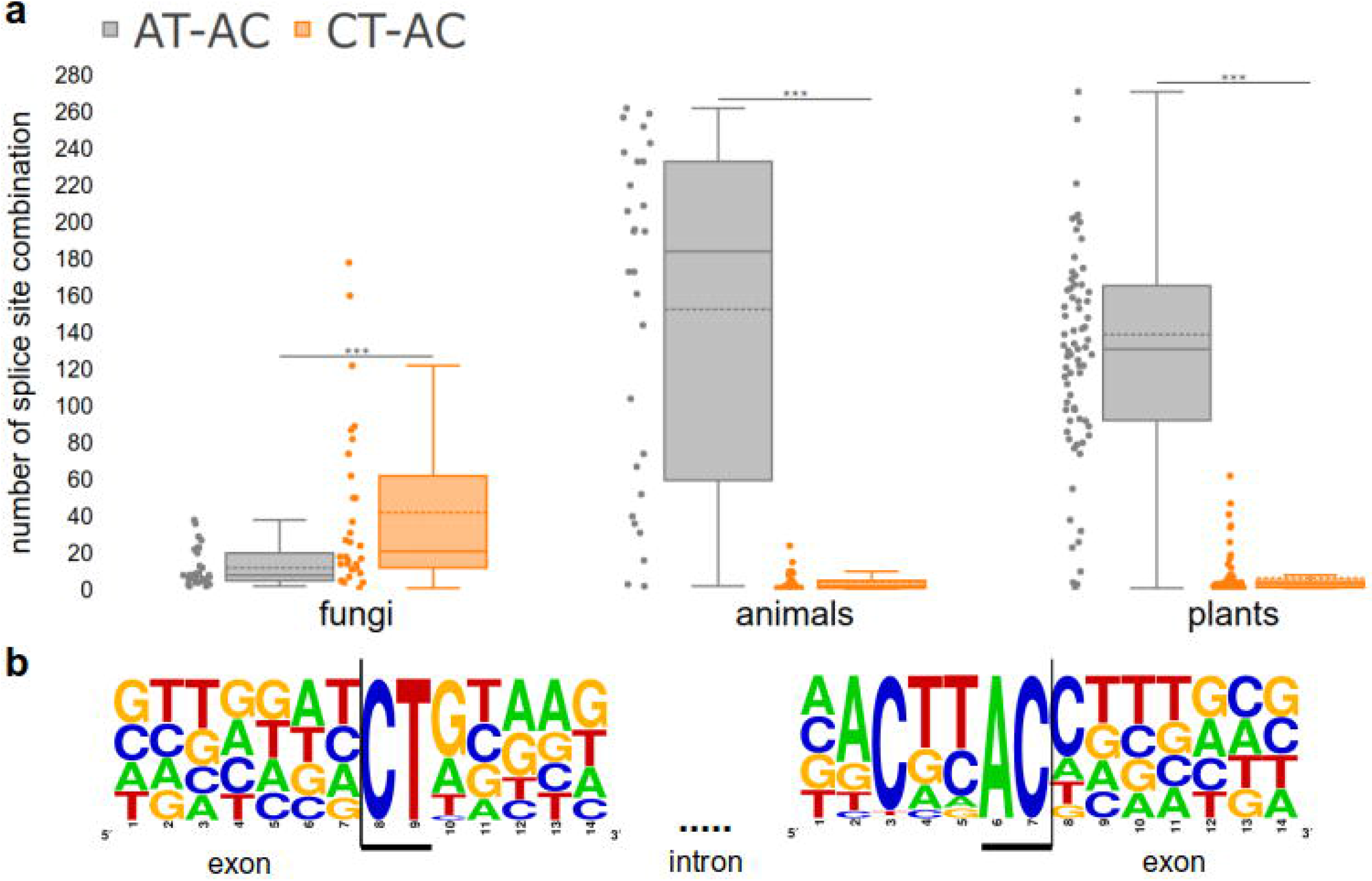
CT-AC frequency exceeds AT-AC frequency in the annotation of fungal genome sequences. A) Number of the minor non-canonical splice site combination CT-AC in comparison to the major non-canonical splice site combination AT-AC in each kingdom (Mann-Whitney U-Test; fungi: p≈0.00035, animals: p≈9.560e-10, plants: p≈5.464e-24). The dashed line represents the mean frequency of the respective splice site combination over all investigated species. B) Sequence logo for the splice site combination CT-AC in four selected fungal species (*Alternaria alternata, Aspergillus brasiliensis, Fomitopsis pinicola* and *Zymoseptoria tritici*). In total, 67 supported splice sites with this combination were used to generate the sequence logo.

The presence of antisense transcripts which would be spliced at a canonical GT-AG splice site combination (reverse complement of CT-AC) is a likely explanation. At least, stranded RNA-Seq data sets are required to investigate this hypothesis in fungi by differentiating between transcripts of both strands. Frequently encountered artifacts caused by reverse transcription^36^ might be avoided in the future through direct RNA sequencing^47^. Due to the very limited availability of suitable data sets for fungal species we have to leave this question for future studies.

### Intron size analysis

Assuming that non-canonical splice sites are not used or used at a lower efficiency, it could be assumed that introns are more often retained than introns with canonical splice sites. A possible consequence of intron retention could be frameshifts unless the intron length is a multiple of three. Therefore, we investigated a total of 8,060,924, 737,783 and 2,785,484 transcripts across animals, fungi, and plants, respectively, with respect to their intron lengths. Introns with a length divisible by three could be kept in the final transcript without causing a shift in the reading frame, because they add complete codons to the transcript. There is no significant difference between introns with different splice site combinations (Table 2). The ratio of introns with a length divisible by 3 is very close to 33.3 % which would be expected based on an equal distribution. The only exception are minor non-canonical splice site combinations in fungi which are slightly less likely to occur in introns with a length divisible by 3.

**Table 2:**
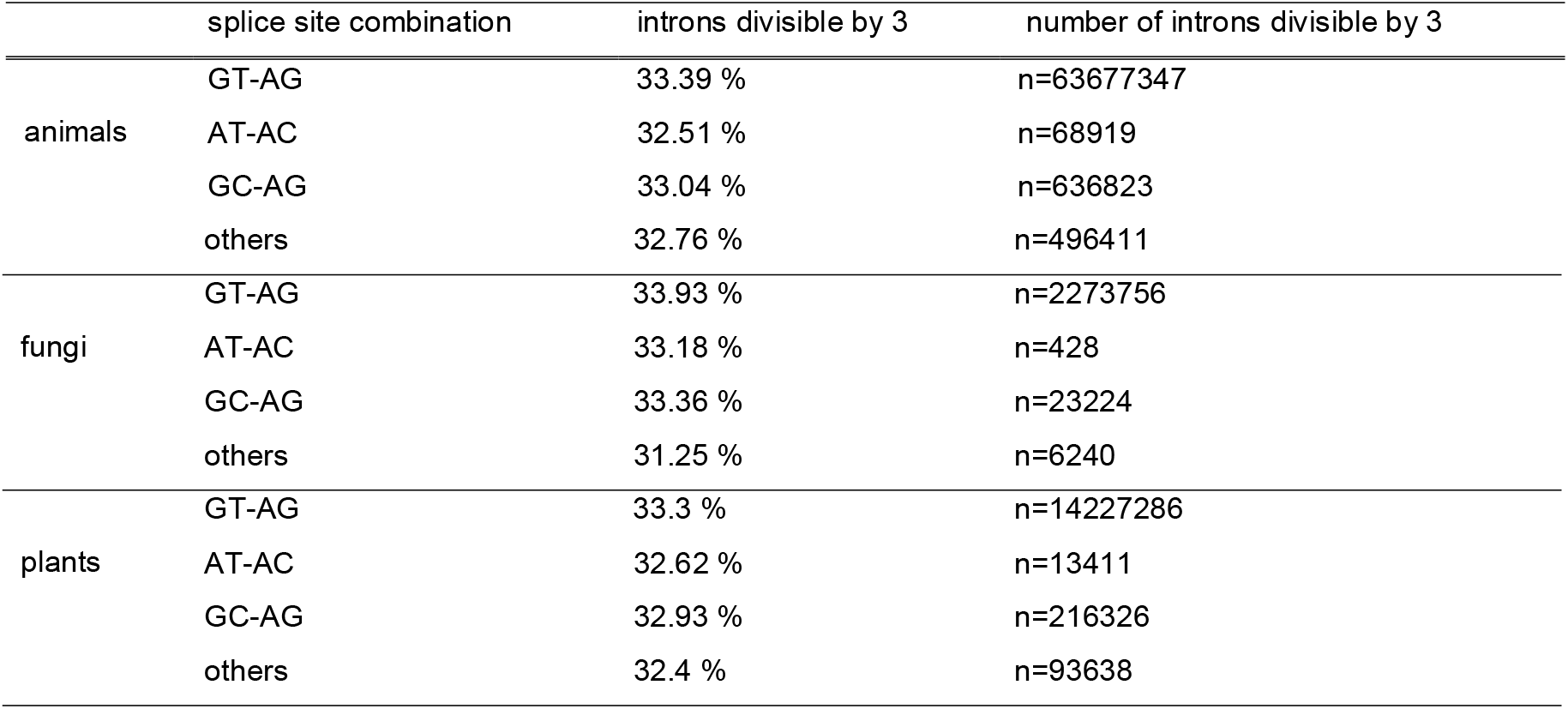
Proportion of introns with length divisible by 3. The results of intron length analysis for selected splice site combinations for animals, fungi and plants are shown.

### Conservation of non-canonical splice site combinations across species

In total, *A. thaliana* transcripts containing 1,073 GC-AG, 64 AT-AC and 19 minor non-canonical splice sites were aligned to transcripts of all plant species. Homologous intron positions were checked for non-canonical splice sites. GC-AG splice site combinations were conserved in 9,830 sequences, matched with other non-canonical splice site combinations in 121 cases, and aligned to GT-AG in 13,045 sequences. Given that the dominance of GT-AG splice sites was around 98 %, the number observed here indicates a strong conservation of GC-AG splice site combinations. AT-AC splice site combinations were conserved in 967 other sequences, matched with other non-canonical splice site combinations in 93 cases, and aligned to GT-AG in 157 sequences. These numbers indicate a conservation of AT-AC splice site combinations, which exceeds the conservation of GC-AG splice site combinations substantially. Minor non-canonical splice sites were conserved in 48 other sequences, matched with other non-canonical splice site combinations in 64 cases, and were aligned to a canonical GT-AG splice site in 213 cases. This pattern suggests that most non-canonical splice site combinations are either (A) mutations of the canonical ones or (B) mutated towards GT-AG splice site combinations.

The power of this analysis is currently limited by the quality of the alignment. Although splice site combinations should be aligned properly in most cases, small differences in the number could be caused by ambiguous situations. It is likely that both events stated above account for a fraction of splice site combinations. To assign each non-canonical splice site combination to A or B, a tool for automatic inspection of the observed phylogenetic pattern would be required. To assess the feasibility of this approach, we investigated the conservation of non-canonical splice sites in transcripts of *Armillaria gallica* as this species shows a high number of non-canonical splice sites in the annotation and the set of fungal genome sequences has a feasible size for this analysis. After identification of putative homologous sequences in other species, phylogenetic trees of these sequences were inspected. Transcripts with non-canonical splice site combinations are clustered in clades which also harbour transcripts without non-canonical splice site combinations. We analysed trees of all transcripts which have similar transcripts with non-canonical splice site combinations in at least 10 other species and observed on average 5 transcripts with a non-canonical splice site combination among the 10 closest relatives. This number is exceeding the expectation based on the overall frequency of less than 3% non-canonical splice site combinations, thus indicating conservation of non-canonical splice sites. Due to this apparently complex evolutionary pattern, we do not know if these clades originated from a non-canonical splice site combination which was turned into a canonical one multiple times (B) or if a non-canonical splice site combination evolved multiple times (A).

### Usage of non-canonical splice sites

Non-canonical splice site combinations were described to have regulatory roles by slowing down the splicing process^48^. Previous reports also indicated that non-canonical splice site combinations might appear in pseudogenes^9,10^. To analyse a possible correlation of non-canonical splice sites with low transcriptional activity, we compared the transcript abundance of genes with non-canonical splice site combinations to genes with only canonical GT-AG splice site combinations (Figure 5A). Genes with at least one non-canonical splice site combination are generally less likely to be lowly expressed than genes with only canonical splice sites. While this trend holds true for all analysed non-canonical splice site combination groups, GC-AG and AT-AC containing genes display especially low proportions of genes with low FPKMs. We speculate that a stronger transcriptional activity of genes with non-canonical splice sites compensates for lower turnover rates in the splicing process. The regulation of these genes might be shifted from the transcriptional to the post-transcriptional level. This trend is similar for animals and plants (Supplementary Data S14). In fungi, genes with minor non-canonical splice sites display relatively high proportions of genes with low FPKMs. Moreover, a higher number of non-canonical splice sites per gene is associated with a lower expression. This leads to the suggestion, that non-canonical splice sites occur more often within pseudogenes.

**Figure 5:**
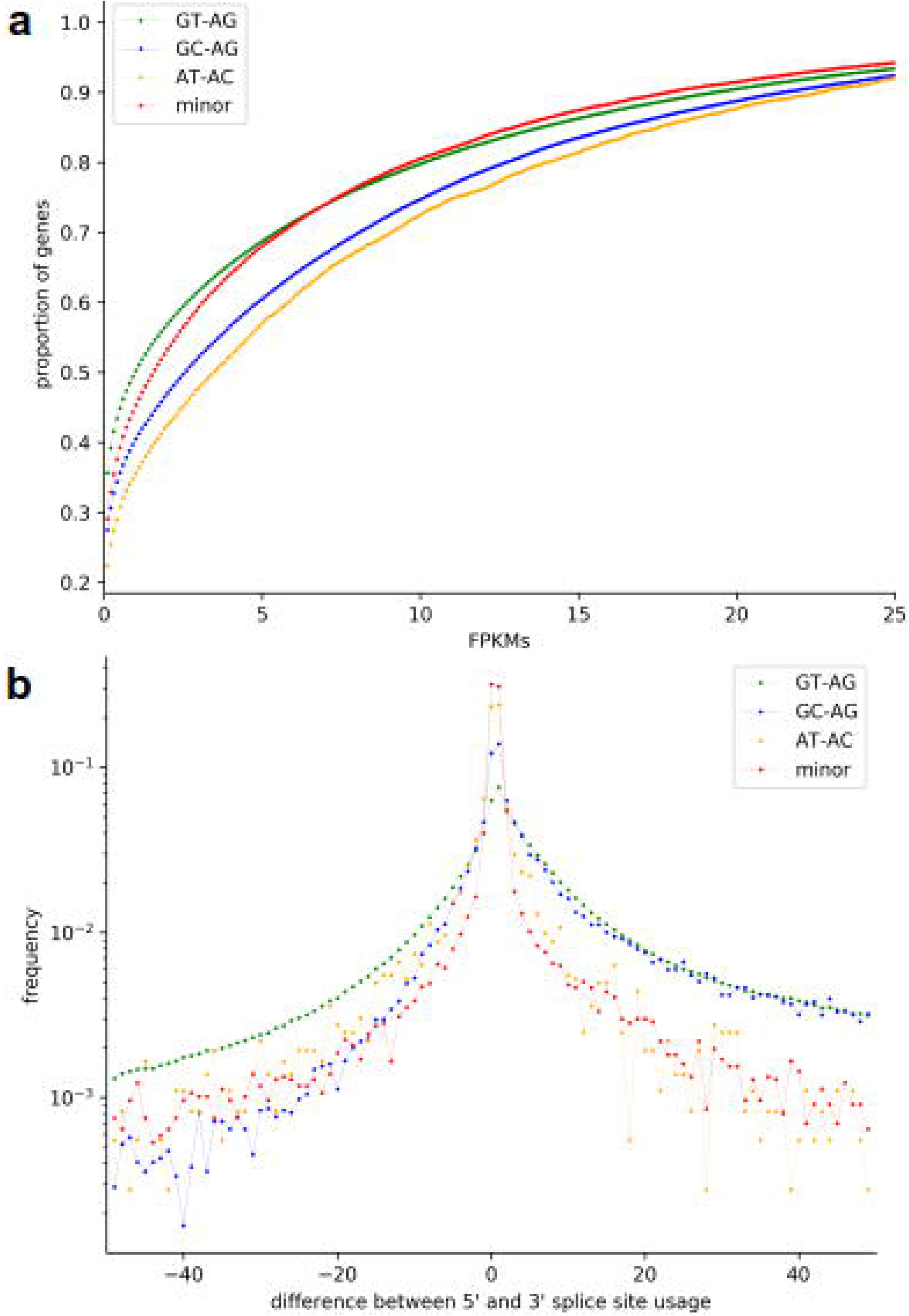
Usage of non-canonical splice sites in plant species. A) Comparison of the transcript abundance (FPKMs) of genes with non-canonical splice site combinations to genes with only canonical GT-AG splice site combinations. GC-AG and AT-AC containing genes display especially low proportions of genes with low FPKMs. This leads to a higher transcript abundance of genes with low FPKMs. B) Comparison of the usage of 5’ and 3’ splice sites. On the x-axis, the difference between the 5’ splice site usage and the usage of the 3’ splice site is shown. A fast drop of values when going to the negative side of the x-axis indicates that the 3’ splice site is probably more flexible than the 5’ splice site.

Introns are mostly defined by phylogenetically conserved splice sites, but nevertheless some variation of these splice sites is possible^9,10,25,26,46^. To understand the amount of flexibility in respect to different terminal dinucleotides, we compared the usage of donor and acceptor splice sites over 4,141,196 introns in plants, 3,915,559 introns in animals, and 340,619 introns in fungi (Figure 5B). The plot shows that the 3’ splice site seems to be more flexible than the 5’ splice site which was observed in all three kingdoms. Our observations align well with previous findings of a higher flexibility at the 3’ splice site compared to the 5’ splice site. A mutated 5’ splice site represses the removal of the upstream intron^10,49,50^. Further, for plants and animals, the difference between the usage of the 5’ splice site and the 3’ splice site is notably higher for introns with the splice site combination GC-AG.

One important limitation of this investigation is the sparse availability of stranded RNA-Seq data sets and direct RNA sequencing data sets. Therefore, it is not possible to rule out the involvement of antisense transcripts, which have been observed before^51^. In three cases, these antisense transcripts could be spliced at a canonical or major non-canonical splice site combination, while appearing as a minor non-canonical splice site on the investigated strand. Previous reports of antisense transcripts^52^ suggest an explanation for this observation. Although *bona fide* non-canonical splice site combinations are present in many plant transcripts^10^, additional transcript isoforms might exist. To evaluate the relevance of such alternative isoforms, we assessed the contribution of isoforms to the overall abundance of transcripts of a gene. Therefore, the usage of splice sites flanking an intron was compared to the average usage of splice sites. This reveals how often a certain intron is removed by splicing. Introns with low usage values might only be involved in minor transcript isoforms. While most introns display no or very small differences, GT-AG introns deviate from this trend. This indicates that non-canonical splice site combinations are frequently part of the dominant isoform. Again, these findings were similar for all of the investigated kingdoms.

Our investigation of non-canonical splice sites in animal, fungal and plant genome sequences revealed kingdom specific differences. Animal species with a high proportion of annotated GA-AG splice site combinations were examined. Further, properties of introns and splice sites were analysed. One aspect of this analysis is, that the 3’ splice site seems to be more flexible than the 5’ splice site, which was observed in all three kingdoms. Across fungal genome sequences, the splice site combination CT-AC is more frequent than the splice site combination AT-AC. This makes CT-AC a major non-canonical splice site combination in fungal species, while AT-AC should be considered a minor non-canonical splice site in fungi. Overall, our findings demonstrate the importance of considering non-canonical splice sites despite their low relative frequency in comparison to the canonical splice site combination GT-AG. RNA-Seq data supported the existence and usage of numerous non-canonical splice site combinations. By neglecting non-canonical splice sites, *bona fide* genes might be excluded or at least structurally altered.

## Methods

### Analysis and validation of splice site combinations

A detailed technical description of all included scripts with usage examples is available from the corresponding github repository (https://github.com/bpucker/ncss2018). A wrapper script is included to automatically perform the analyses based on provided genome sequence (FASTA), annotation (GFF3), and RNA-Seq read mapping (BAM) per species of interest.

Genome sequences (FASTA) and corresponding annotations (GFF3) of 130 fungal species and 489 animal species were retrieved from the NCBI. Representative transcript and peptide sequences were extracted as described before^10^. General statistics were calculated using a Python script^10^. The completeness of all data sets was assessed with BUSCO v3^53^ running in protein mode on the representative peptide sequences using the reference data sets ‘fungi odb9’ and ‘metazoa odb9’, respectively^54^ (Supplementary Data S15 and Supplementary Data S16).

To validate the detected splice site combinations, paired-end RNA-Seq data sets were retrieved from the Sequence Read Archive^55^ (Supplementary Data S17 and Supplementary Data S18). The following validation approach^10^ utilized STAR v2.5.1b^56^ for the read mapping and Python scripts for downstream processing (https://doi.org/10.5281/zenodo.2586989). RNA-Seq reads were considered mapped if the alignment shows at least 95% similarity and covers 90% of the read length. Splice sites were considered valid if they are spanned by at least three reads and show a coverage drop of 20% when moving from an exon into an intron sequence. Summaries of the RNA-Seq read coverage depth at splice sites in animals^57^ and fungi^58^ have been made available as part of this study.

RNA-Seq read mappings with STAR v2.5.1b and HiSat2 v.2.1.0 were compared based on a gold standard generated by exonerate, because a previous report^59^ indicated a superiority of STAR. STAR parameters were set as described above and HiSat2 was applied with default parameters. All transcripts with non-canonical splice sites in *A. thaliana* and *Oryza sativa* were considered. When investigating the alignment of RNA-Seq reads over non-canonical splice sites, we observed a high accuracy for both mappers without a clear difference between them. Previously described scripts^10^ were adjusted for this analysis and updated versions are available on github (https://doi.org/10.5281/zenodo.2586989). The distribution of genome sizes was analysed using the Python package dabest^60^. Sequence logos for the analysed splice sites were designed at http://weblogo.berkeley.edu/logo.cgi^61^.

### Calculation of the splice site diversity

A custom Python script (splice_site_divergence_check.py) was applied to calculate the Shannon diversity index (H’)^62^ of all splice site combinations in fungi, animals and plants (https://doi.org/10.5281/zenodo.2586989). To determine the significance of the obtained results, a Kruskal-Wallis test^63^ was calculated using the Python package scipy^64^. Further, the interquartile range of all distributions was examined.

### Investigation of a common non-canonical splice site in fungal genome sequences

A Mann-Whitney U Test implemented in the Python package scipy was performed to analyse differences in the number of minor non-canonical splice site combinations. The observed distributions were visualized in a boxplot (https://doi.org/10.5281/zenodo.2586989) constructed with the Python package plotly^65^ (ss_combination_frequency_boxplot.py).

### Detection of spliceosomal components

A potential U1 snRNA of *Pan troglodytes* (obtained from the NCBI; GeneID:112207549) was subjected to BLASTn v.2.8.1^66^ against the genome sequences of selected species. Hits with a score above 100, with at least 80 % similarity and with the conserved sequence at the 5’ end of the snRNA^67^ were investigated, as these sequences are potential U1 snRNAs. The obtained sequences were compared and small nucleotide variants were detected.

To assess possible duplications of spliceosomal components, bait sequences from various species were collected for central proteins in the spliceosome including pre-mRNA-processing factor 8 (PRP8), U1 small nuclear ribonucleoprotein C, U4/U6 small nuclear ribonucleoprotein Prp3, U4/U6.U5 small nuclear ribonucleoprotein 27 kDa protein and U5 small nuclear ribonucleoprotein. Putative homologues in all animal species were detected based on Python scripts^68^ and subjected to the construction of phylogenetic trees as described previously^69^.

Genome sequences were systematically screened for U12 spliceosome hints via Infernal (cmscan) v1.1.2^70^ with Rfam13^71^. U4atac, U6atac, U11, and U12 were considered as indications for the presence of the minor U12 spliceosome in the respective species. Due to high computational costs only a random subset of all animal sequences was analysed.

### Correlation between the GC content of the genome and the GC content of the splice sites

The Pearson correlation coefficient between the GC content of the genome sequence of each species and the GC content of the respective splice site combination was calculated using the Python package scipy. Splice site combinations were weighted with the number of occurrences for assessment of the GC content. Finally, the correlation coefficient and the p-value were determined. For better visualization, a scatter plot was constructed with the Python package plotly^65^.

### Phylogeny of non-canonical splice sites

All *A. thaliana* transcripts with non-canonical splice sites were subjected to BLASTn v.2.8.1 searches against the transcript sequences of all other plant species previously studied^10^. The best hit per species was selected for an alignment against the respective genomic region with exonerate^72^. Next, splice site combinations were extracted and aligned. This alignment utilized MAFFT v7^73^ by representing different splice site combinations as amino acids. Finally, splice site combinations aligned with the non-canonical splice site combinations of *A. thaliana* were analysed (https://doi.org/10.5281/zenodo.2586989).

All transcripts of the fungus *Armillaria gallica* with non-canonical splice sites were searched as translated peptide sequences against all other fungal peptide sequence data sets via BLASTp^66^. Cases with more than 10 best hits with non-canonical splice site combinations in other species were subjected to the construction of phylogenetic trees for manual inspection. MAFFT v7^73^ and FastTree v2^74^ were used for the alignment and tree construction.

### Usage of non-canonical splice sites

Genes were classified based on the presence/absence of non-canonical splice combinations into four groups: GT-AG, GC-AG, AT-AC, and minor non-canonical splice site genes. When having different non-canonical splice sites, genes were assigned into multiple groups. Next, the transcription of these genes was quantified based on RNA-Seq using featureCounts v1.5.0-p3^75^ based on the RNA-Seq read mapping generated with STAR v.2.5.1b. Multi-mapped reads were excluded from the analysis and expression values were calculated at gene level (-t gene). Binning of the genes was performed based on the fragments per kilobase transcript length per million assigned fragments (FPKMs). Despite various shortcomings^76^, we consider FPKMs to be acceptable for this analysis. Outlier genes with extremely high values were excluded from this analysis and the visualization. Next, a cumulative sum of the relative bin sizes was calculated. The aim was to compare the transcriptional activity of genes with different splice site combinations i.e. to test whether non-canonical splice site combinations are enriched in lowly transcribed genes.

Usage of splice sites was calculated per intron as previously described^10^. The difference between both ends of an intron was calculated. The distribution of these differences per splice site type were analysed.

Introns were grouped by their splice site combination. The average of both coverage values of the directly flanking exon positions was calculated as estimate of the local expression around a splice site combination. Next, the sequencing coverage of a transcript was estimated by multiplying 200 bp (assuming 2×100 nt reads) with the number of read counts per gene and normalization to the transcript length. The difference between both values was calculated for each intron to assess its presence in the major isoform.

### Genomic read mapping and variant calling

Genomic sequencing reads were retrieved from the SRA via fastq-dump as described above. BWA MEM v.0.7^77^ was applied with the -m parameter for mapping of the reads and GATK v.3.8^78,79^ was used for the variant detection as described previously^80^. The positions of variants were compared to the positions of splice sites using compare_variation_rates.py (https://github.com/bpucker/ncss2018).

### Data Availability

This work was based on publicly available data sets retrieved from the NCBI (Supplementary Data S3, Supplementary Data S4) and the SRA (Supplementary Data S17, Supplementary Data S18). Python scripts and a detailed technical description are available at github: https://github.com/bpucker/ncss2018. Data sets with information about the coverage around splice sites in animals (http://doi.org/10.4119/unibi/2934226) and fungi (http://doi.org/10.4119/unibi/2934220) were made available as data publication at Bielefeld University Library.

## Acknowledgments

We thank members of Genetics and Genomics of Plants for discussion of preliminary results. We are very grateful to Hanna Marie Schilbert, Janik Sielemann, and Iain Place for helpful comments on the manuscript. We acknowledge support for the Article Processing Charge by the Deutsche Forschungsgemeinschaft and the Open Access Publication Fund of Bielefeld University.

## Author Contributions Statement

K.F. and B.P. designed the study, performed the experiments, analysed the data, and wrote the manuscript. Both authors read and approved the final version of this manuscript.

## Competing interests

The authors declare no competing interests.

## Supplementary Data

SupplementaryData S**1.** List of all possible splice site combinations in animal species.

Supplementary Data S**2.** List of all possible splice site combinations in fungal species.

Supplementary Data S**3.** List of genome sequences and annotations of the investigated animal species.

Supplementary S**4.** List of genome sequences and annotations of the investigated fungal species.

Supplementary Data S**5.** Distribution of genome sizes of all species.

Supplementary Data S**6.** Genome statistics concerning each analysed animal species and status of U12 spliceosome.

Supplementary Data S**7.** Genome statistics concerning each analysed fungal species and status of U12 spliceosome.

Supplementary Data S**8.** Distribution of canonical and non-canonical splice sites per species in the animal kingdom.

Supplementary Data S**9.** Distribution of canonical and non-canonical splice sites per species in the fungal kingdom.

Supplementary Data S**10.** Percentage of variants in canonical splice sites, major and minor non-canonical splice sites.

Supplementary Data S**11.** Correlation between the GC content of the genome and the GC content of the splice sites per kingdom.

Supplementary Data S**12.** Correlation between the size of the used RNA-Seq data sets and the number of supported splice sites.

Supplementary Data S**13.** Proportion of genes with non-canonical splice sites in dependence of the number of introns.

Supplementary Data S**14.** Usage of non-canonical splice sites in animals and fungi.

Supplementary Data S**15.** Non-canonical splice sites in BUSCOs and in all genes were assessed per species in the animal kingdom.

Supplementary Data S**16.** Non-canonical splice sites in BUSCOs and in all genes were assessed per species in the fungal kingdom.

Supplementary Data S**17.** List of Sequence Read Archive accession numbers of the investigated animal RNA-Seq data sets.

Supplementary Data S**18.** List of Sequence Read Archive accession numbers of the investigated fungal RNA-Seq data sets.

## Notes

#### Summary of Updates

Substantially improved version of the manuscript. New Figure added.

https://github.com/bpucker/ncss2018

http://doi.org/10.4119/unibi/2934220

http://doi.org/10.4119/unibi/2934226

## References

1. Moore, Melissa J and Sharp, Phillip A. Site-specific modification of pre-mRNA: the 2’-hydroxyl groups at the splice sites. Science 256, 992–997 (1992).

2. Barbosa-Morais, Nuno L and Irimia, Manuel and Pan, Qun and Xiong, Hui Y and Gueroussov, Serge and Lee, Leo J and Slobodeniuc, Valentina and Kutter, Claudia and Watt, Stephen and Çolak, Recep and others. The evolutionary landscape of alternative splicing in vertebrate species. Science 338, 1587–1593 (2012).

3. Ben-Dov, Claudia and Hartmann, Britta and Lundgren, Josefin and Valcárcel, Juan. Genome-wide analysis of alternative pre-mRNA splicing. J. Biol. Chem. 283, 1229–1233 (2008).

4. Matlin, Arianne J and Clark, Francis and Smith, Christopher WJ. Understanding alternative splicing: towards a cellular code. Nat. Rev. Mol. Cell Biol. 6, 386 (2005).

5. Sibley, Christopher R and Blazquez, Lorea and Ule, Jernej. Lessons from non-canonical splicing. Nat. Rev. Genet. 17, 407 (2016).

6. Maniatis, Tom and Tasic, Bosiljka. Alternative pre-mRNA splicing and proteome expansion in metazoans. Nature 418, 236 (2002).

7. Xue, Min and Chen, Bing and Ye, Qingqing and Shao, Jingru and Lyu, Zhangxia and Wen, Jianfan. Sense-antisense gene overlap causes evolutionary retention of the few introns in Giardia genome and the implications. bioRxiv (2018).

8. Chorev, Michal and Carmel, Liran. The function of introns. Front. Genet. 3, (2012).

9. Burset, M and Seledtsov, IA and Solovyev, VV. Analysis of canonical and non-canonical splice sites in mammalian genomes. Nucleic Acids Res. 28, 4364–4375 (2000).

10. Pucker, Boas and Brockington, Samuel F. Genome-wide analyses supported by RNA-Seq reveal non-canonical splice sites in plant genomes. BMC Genomics 19, 980 (2018).

11. Bon, Elisabeth and Casaregola, Serge and Blandin, Gaëlle and Llorente, Bertrand and Neuvéglise, Cécile and Munsterkotter, Martin and Guldener, Ulrich and Mewes, Hans-Werner and Helden, Jacques Van and Dujon, Bernard and others. Molecular evolution of eukaryotic genomes: hemiascomycetous yeast spliceosomal introns. Nucleic Acids Res. 31, 1121–1135 (2003).

12. Logsdon, John M. The recent origins of spliceosomal introns revisited. Curr. Opin. Genet. Dev. 8, 637–648 (1998).

13. Burge, Chris and Karlin, Samuel. Prediction of complete gene structures in human genomic DNA1. J. Mol. Biol. 268, 78–94 (1997).

14. Stanke, Mario and Waack, Stephan. Gene prediction with a hidden Markov model and a new intron submodel. Bioinformatics 19, ii215–ii225 (2003).

15. Davis, Carrie A and Grate, Leslie and Spingola, Marc and Ares Jr, Manuel. Test of intron predictions reveals novel splice sites, alternatively spliced mRNAs and new introns in meiotically regulated genes of yeast. Nucleic Acids Res. 28, 1700–1706 (2000).

16. Wahl, Markus C and Will, Cindy L and Lührmann, Reinhard. The spliceosome: design principles of a dynamic RNP machine. Cell 136, 701–718 (2009).

17. Sharp, Phillip A and Burge, Christopher B. Classification of introns: U2-type or U12-type. Cell 91, 875–879 (1997).

18. Hall, Stephen L and Padgett, Richard A. Requirement of U12 snRNA for in vivo splicing of a minor class of eukaryotic nuclear pre-mRNA introns. Science 271, 1716–1718 (1996).

19. Turunen, Janne J and Niemelä, Elina H and Verma, Bhupendra and Frilander, Mikko J. The significant other: splicing by the minor spliceosome. Wiley Interdiscip. Rev. RNA 4, 61–76 (2013).

20. Dietrich, Rosemary C and Incorvaia, Robert and Padgett, Richard A. Terminal intron dinucleotide sequences do not distinguish between U2-and U12-dependent introns. Mol. Cell 1, 151–160 (1997).

21. Wilkinson, Max E and Fica, Sebastian M and Galej, Wojciech P and Norman, Christine M and Newman, Andrew J and Nagai, Kiyoshi. Postcatalytic spliceosome structure reveals mechanism of 3’–splice site selection. Science 358, 1283–1288 (2017).

22. Burge, Christopher B and Tuschl, Thomas and Sharp, Phillip A. Splicing of precursors to mRNAs by the spliceosomes. Cold Spring Harb. Monogr. Ser. 37, 525–560 (1999).

23. Roca, Xavier and Krainer, Adrian R and Eperon, Ian C. Pick one, but be quick: 5’ splice sites and the problems of too many choices. Genes Dev. 27, 129–144 (2013).

24. Shi, Yigong. The spliceosome: a protein-directed metalloribozyme. J. Mol. Biol. 429, 2640–2653 (2017).

25. Kupfer, Doris M and Drabenstot, Scott D and Buchanan, Kent L and Lai, Hongshing and Zhu, Hua and Dyer, David W and Roe, Bruce A and Murphy, Juneann W. Introns and splicing elements of five diverse fungi. Eukaryot. Cell 3, 1088–1100 (2004).

26. Kitamura-Abe, Sumie and Itoh, Hitomi and Washio, Takanori and Tsutsumi, Akihiro and Tomita, Masaru. Characterization of the splice sites in GT–AG and GC–AG introns in higher eukaryotes using full-length cDNAs. J. Bioinform. Comput. Biol. 2, 309–331 (2004).

27. Michael, Deutsch and Manyuan, Long. Intron—exon structures of eukaryotic model organisms. Nucleic Acids Res. 27, 3219–3228 (1999).

28. Modrek, Barmak and Resch, Alissa and Grasso, Catherine and Lee, Christopher. Genome-wide detection of alternative splicing in expressed sequences of human genes. Nucleic Acids Res. 29, 2850–2859 (2001).

29. Pucker, Boas and Holtgräwe, Daniela and Weisshaar, Bernd. Consideration of non-canonical splice sites improves gene prediction on the Arabidopsis thaliana Niederzenz-1 genome sequence. BMC Res. Notes 10, 667 (2017).

30. Sparks, Michael E and Brendel, Volker. Incorporation of splice site probability models for non-canonical introns improves gene structure prediction in plants. Bioinformatics 21, iii20–iii30 (2005).

31. Dubrovina, AS and Kiselev, KV and Zhuravlev, Yu N. The role of canonical and noncanonical pre-mRNA splicing in plant stress responses. BioMed Res. Int. 2013, (2013).

32. Alexandrov, Nickolai N and Troukhan, Maxim E and Brover, Vyacheslav V and Tatarinova, Tatiana and Flavell, Richard B and Feldmann, Kenneth A. Features of Arabidopsis genes and genome discovered using full-length cDNAs. Plant Mol. Biol. 60, 69–85 (2006).

33. Niu, Xiangli and Luo, Di and Gao, Shaopei and Ren, Guangjun and Chang, Lijuan and Zhou, Yuke and Luo, Xiaoli and Li, Yuxiang and Hou, Pei and Tang, Wei and others. A conserved unusual posttranscriptional processing mediated by short, direct repeated (SDR) sequences in plants. J. Genet. Genomics 37, 85–99 (2010).

34. Erkelenz, Steffen and Theiss, Stephan and Kaisers, Wolfgang and Ptok, Johannes and Walotka, Lara and Müller, Lisa and Hillebrand, Frank and Brillen, Anna-Lena and Sladek, Michael and Schaal, Heiner. Ranking noncanonical 5’ splice site usage by genome-wide RNA-seq analysis and splicing reporter assays. Genome Res. 28, 1826–1840 (2018).

35. Grützmann, Konrad and Szafranski, Karol and Pohl, Martin and Voigt, Kerstin and Petzold, Andreas and Schuster, Stefan. Fungal alternative splicing is associated with multicellular complexity and virulence: a genome-wide multi-species study. DNA Res. 21, 27–39 (2013).

36. Parada, G. E., Munita, R., Cerda, C. A. & Gysling, K. A comprehensive survey of non-canonical splice sites in the human transcriptome. Nucleic Acids Res. 42, 10564–10578 (2014).

37. Keren, H., Lev-Maor, G. & Ast, G. Alternative splicing and evolution: diversification, exon definition and function. Nat. Rev. Genet. 11, 345 (2010).

38. Jackson, I. J. A reappraisal of non-consensus mRNA splice sites. Nucleic Acids Res. 19, 3795 (1991).

39. Robertson, Hugh M. Non-canonical GA and GG 5’Intron Donor Splice Sites Are Common in the Copepod Eurytemora affinis. G3 Genes Genomes Genet. g3–300189 (2017).

40. Lee, C. E. Evolutionary mechanisms of habitat invasions, using the copepod Eurytemora affinis as a model system. Evol. Appl. 9, 248–270 (2016).

41. Seo, H.-C. et al. Miniature genome in the marine chordate Oikopleura dioica. Science 294, 2506–2506 (2001).

42. Brackenridge, Simon and Wilkie, Andrew OM and Screaton, Gavin R. Efficient use of a ‘dead-end’GA 5” splice site in the human fibroblast growth factor receptor genes’. EMBO J. 22, 1620–1631 (2003).

43. Mount, S. M. A catalogue of splice junction sequences. Nucleic Acids Res. 10, 459–472 (1982).

44. Tyler, Brett M. Phytophthora sojae: root rot pathogen of soybean and model oomycete. Mol. Plant Pathol. 8, 1–8 (2007).

45. Förster, Helga and Coffey, Michael D and Elwood, Hille and Sogin, Mitchell L. Sequence analysis of the small subunit ribosomal RNAs of three zoosporic fungi and implications for fungal evolution. Mycologia 306–312 (1990).

46. Shen, Danyu and Ye, Wenwu and Dong, Suomeng and Wang, Yuanchao and Dou, Daolong. Characterization of intronic structures and alternative splicing in Phytophthora sojae by comparative analysis of expressed sequence tags and genomic sequences. Can. J. Microbiol. 57, 84–90 (2011).

47. Garalde, D. R. et al. Highly parallel direct RNA sequencing on an array of nanopores. Nat. Methods 15, 201 (2018).

48. Aebi, M and Hornig, H and Padgett, RA and Reiser, J and Weissmann, C. Sequence requirements for splicing of higher eukaryotic nuclear pre-mRNA. Cell 47, 555–565 (1986).

49. Talerico, MELISSA and Berget, SUSAN M. Effect of 5’splice site mutations on splicing of the preceding intron. Mol. Cell. Biol. 10, 6299–6305 (1990).

50. Berget, Susan M. Exon recognition in vertebrate splicing. J. Biol. Chem. 270, 2411–2414 (1995).

51. Kuhn, J. M., Breton, G. & Schroeder, J. I. mRNA metabolism of flowering-time regulators in wild-type Arabidopsis revealed by a nuclear cap binding protein mutant, abh1. Plant J. 50, 1049–1062 (2007).

52. Donaldson, M. E. & Saville, B. J. Natural antisense transcripts in fungi: Natural antisense transcripts in fungi. Mol. Microbiol. 85, 405–417 (2012).

53. Simão, Felipe A and Waterhouse, Robert M and Ioannidis, Panagiotis and Kriventseva, Evgenia V and Zdobnov, Evgeny M. BUSCO: assessing genome assembly and annotation completeness with single-copy orthologs. Bioinformatics 31, 3210–3212 (2015).

54. Kriventseva, Evgenia V and Tegenfeldt, Fredrik and Petty, Tom J and Waterhouse, Robert M and Simao, Felipe A and Pozdnyakov, Igor A and Ioannidis, Panagiotis and Zdobnov, Evgeny M. OrthoDB v8: update of the hierarchical catalog of orthologs and the underlying free software. Nucleic Acids Res. 43, D250–D256 (2014).

55. Leinonen, Rasko and Sugawara, Hideaki and Shumway, Martin and International Nucleotide Sequence Database Collaboration. The sequence read archive. Nucleic Acids Res. 39, D19–D21 (2010).

56. Dobin, Alexander and Davis, Carrie A and Schlesinger, Felix and Drenkow, Jorg and Zaleski, Chris and Jha, Sonali and Batut, Philippe and Chaisson, Mark and Gingeras, Thomas R. STAR: ultrafast universal RNA-seq aligner. Bioinformatics 29, 15–21 (2013).

57. Pucker, Boas and Frey, Katharina. RNA-Seq read coverage depth of splice sites in animals. (2019).

58. Pucker, Boas and Frey, Katharina. RNA-Seq read coverage depth of splice sites in fungi. (2019).

59. Dobin, Alexander and Gingeras, Thomas R. Comment on “TopHat2: accurate alignment of transcriptomes in the presence of insertions, deletions and gene fusions” by Kim et al. https://www.biorxiv.org/content/10.1101/000851v1.full?%3Fcollection= (2013).

60. Ho, Joses and Tumkaya, Tayfun and Aryal, Sameer and Choi, Hyungwon and Claridge-Chang, Adam. Moving beyond P values: Everyday data analysis with estimation plots. bioRxiv 377978 (2018).

61. Crooks, Gavin E and Hon, Gary and Chandonia, John-Marc and Brenner, Steven E. WebLogo: a sequence logo generator. Genome Res. 14, 1188–1190 (2004).

62. Heip, Carlo. A new index measuring evenness. J. Mar. Biol. Assoc. U. K. 54, 555–557 (1974).

63. Breslow, Norman. A generalized Kruskal-Wallis test for comparing K samples subject to unequal patterns of censorship. Biometrika 57, 579–594 (1970).

64. Eric Jones and Travis Oliphant and Pearu Peterson and others. SciPy: Open source scientific tools for Python. (2001).

65. Plotly Technologies Inc. Collaborative data science. https://plot.ly (2015).

66. Altschul, Stephen F and Gish, Warren and Miller, Webb and Myers, Eugene W and Lipman, David J. Basic local alignment search tool. J. Mol. Biol. 215, 403–410 (1990).

67. Stark, Holger and Dube, Prakash and Lührmann, Reinhard and Kastner, Berthold. Arrangement of RNA and proteins in the spliceosomal U1 small nuclear ribonucleoprotein particle. Nature 409, 539 (2001).

68. Yang, Y. et al. Dissecting Molecular Evolution in the Highly Diverse Plant Clade Caryophyllales Using Transcriptome Sequencing. Mol. Biol. Evol. 32, 2001–2014 (2015).

69. Schilbert, H. M., Pellegrinelli, V., Rodriguez-Cuenca, S., Vidal-Puig, A. & Pucker, B. Harnessing natural diversity to identify key amino acid residues in prolidase. http://biorxiv.org/lookup/doi/10.1101/423475 (2018) doi:10.1101/423475.

70. Nawrocki, E. P. & Eddy, S. R. Infernal 1.1: 100-fold faster RNA homology searches. Bioinformatics 29, 2933–2935 (2013).

71. Kalvari, I. et al. Rfam 13.0: shifting to a genome-centric resource for noncoding RNA families. Nucleic Acids Res. 46, D335–D342 (2018).

72. Slater, Guy St C and Birney, Ewan. Automated generation of heuristics for biological sequence comparison. BMC Bioinformatics 6, 31 (2005).

73. Katoh, Kazutaka and Standley, Daron M. MAFFT multiple sequence alignment software version 7: improvements in performance and usability. Mol. Biol. Evol. 30, 772–780 (2013).

74. Price, Morgan N and Dehal, Paramvir S and Arkin, Adam P. FastTree 2–approximately maximum-likelihood trees for large alignments. PloS One 5, e9490 (2010).

75. Liao, Yang and Smyth, Gordon K and Shi, Wei. featureCounts: an efficient general purpose program for assigning sequence reads to genomic features. Bioinformatics 30, 923–930 (2013).

76. Conesa, Ana and Madrigal, Pedro and Tarazona, Sonia and Gomez-Cabrero, David and Cervera, Alejandra and McPherson, Andrew and Szcześniak, Micha\l Wojciech and Gaffney, Daniel J and Elo, Laura L and Zhang, Xuegong and others. A survey of best practices for RNA-seq data analysis. Genome Biol. 17, 13 (2016).

77. Li, H. Aligning sequence reads, clone sequences and assembly contigs with BWA-MEM. ArXiv Prepr. ArXiv13033997 (2013).

78. McKenna, A. et al. The Genome Analysis Toolkit: a MapReduce framework for analyzing next-generation DNA sequencing data. Genome Res. 20, 1297–1303 (2010).

79. Van der Auwera, G. A. et al. From FastQ data to high-confidence variant calls: the genome analysis toolkit best practices pipeline. Curr. Protoc. Bioinforma. 43, 11–10 (2013).

80. Baasner, J.-S., Howard, D. & Pucker, B. Influence of neighboring small sequence variants on functional impact prediction. BioRxiv (2019).

