## Supplementary figures and images for "Animal, fungi, and plant genome sequences harbour different non-canonical splice sites"

### Supplementary Data S5

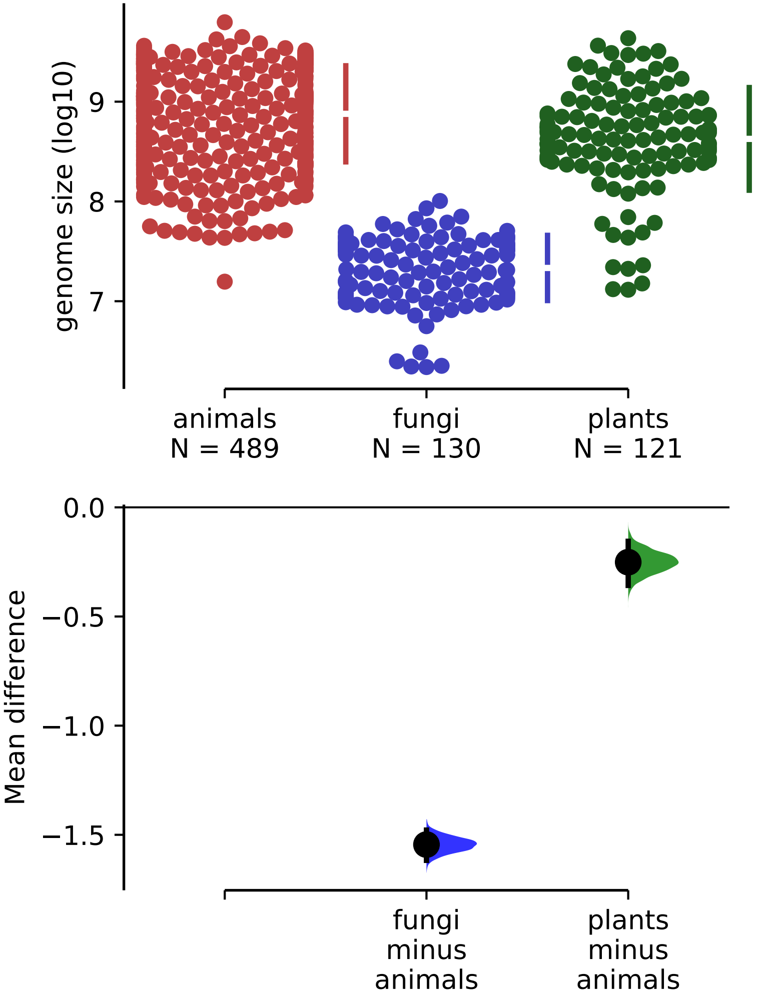

### Supplementary Data S11

#### animals

r = 0.43437813971655936, p = 7.866123362747553e-24

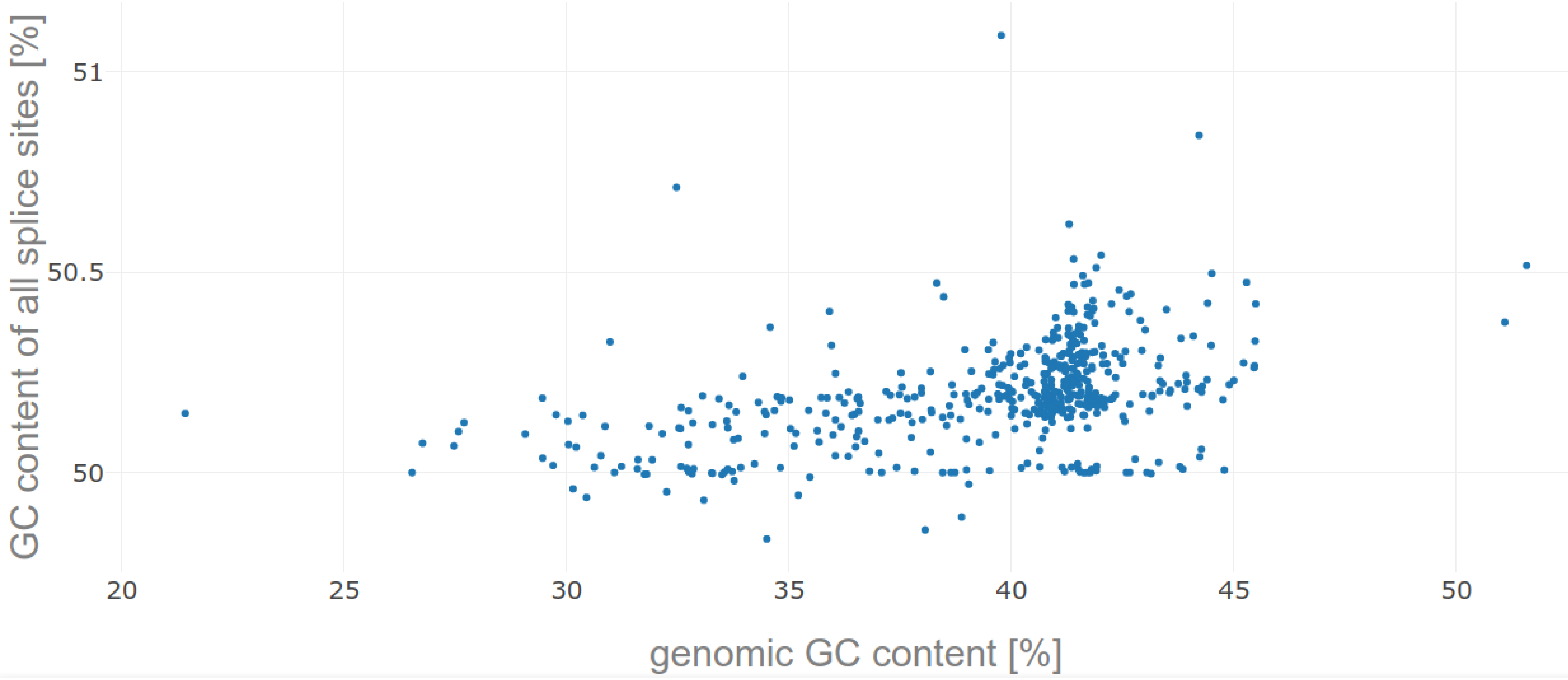

#### fungi

r = 0.23568093817494543, p = 0.007891452637704533

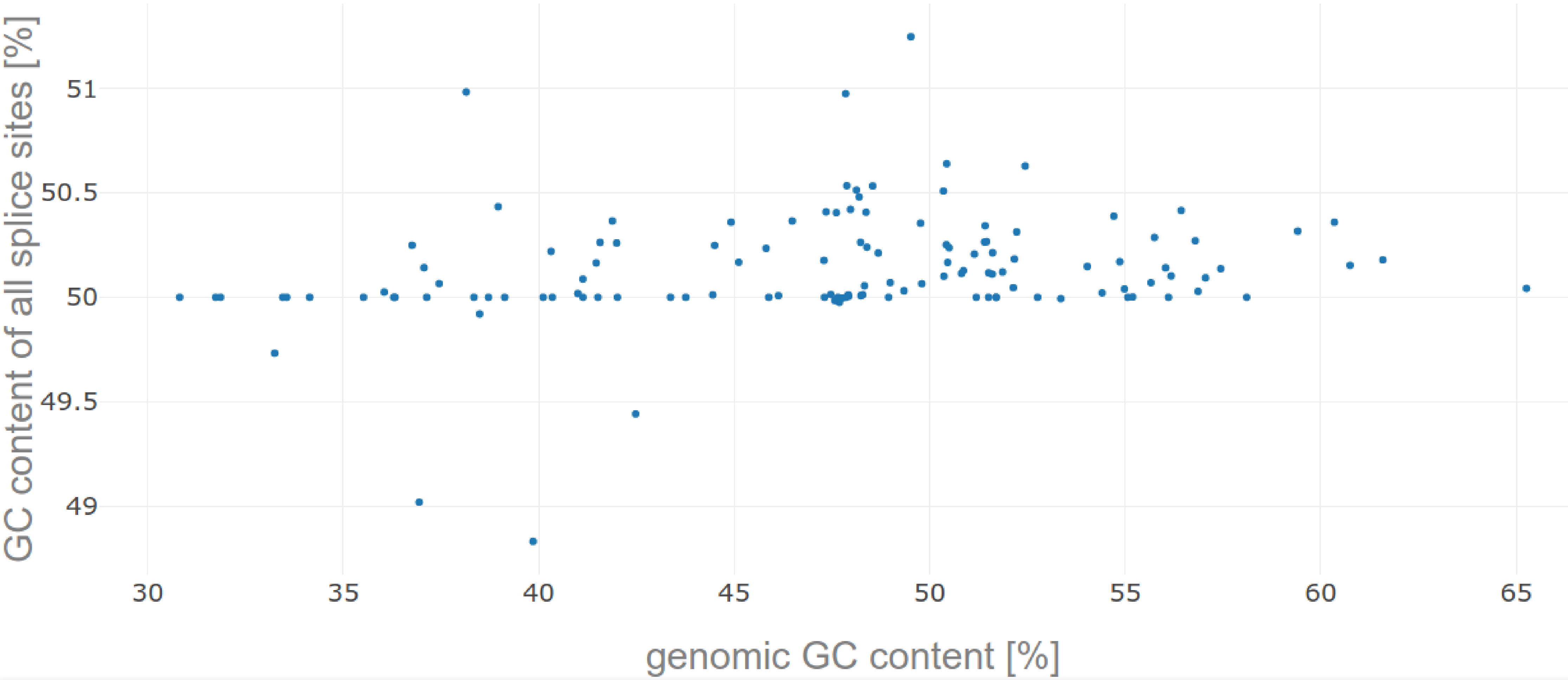

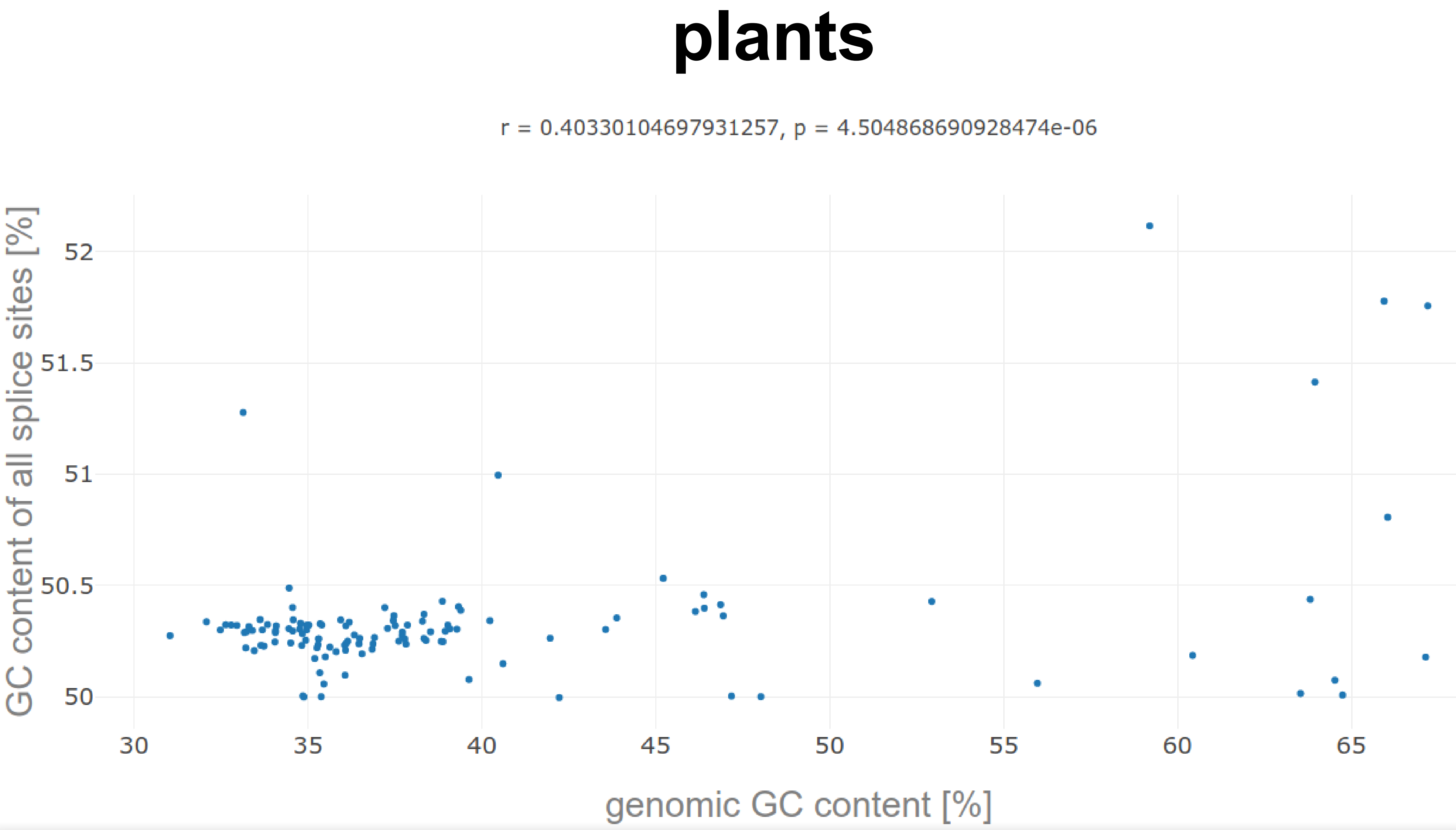

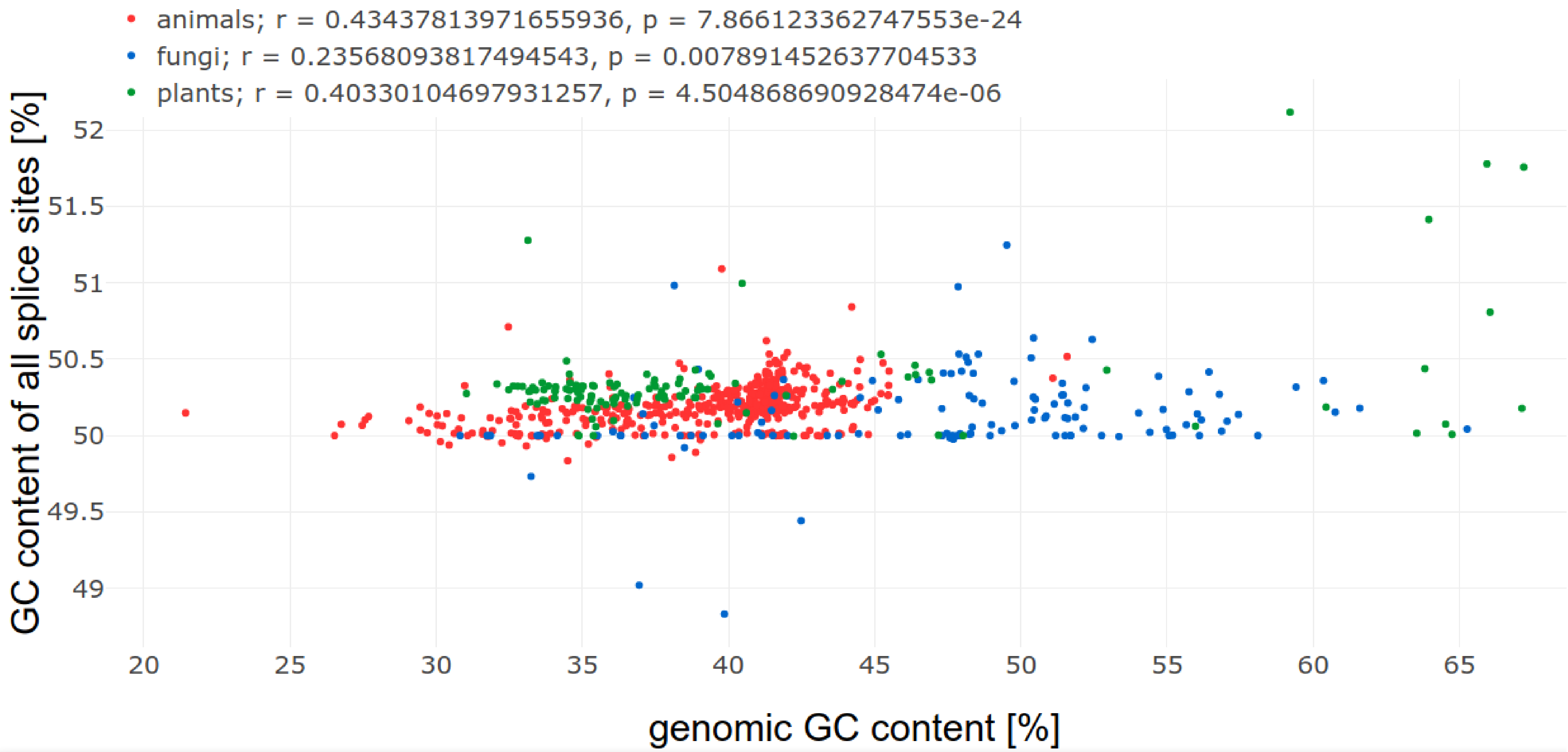

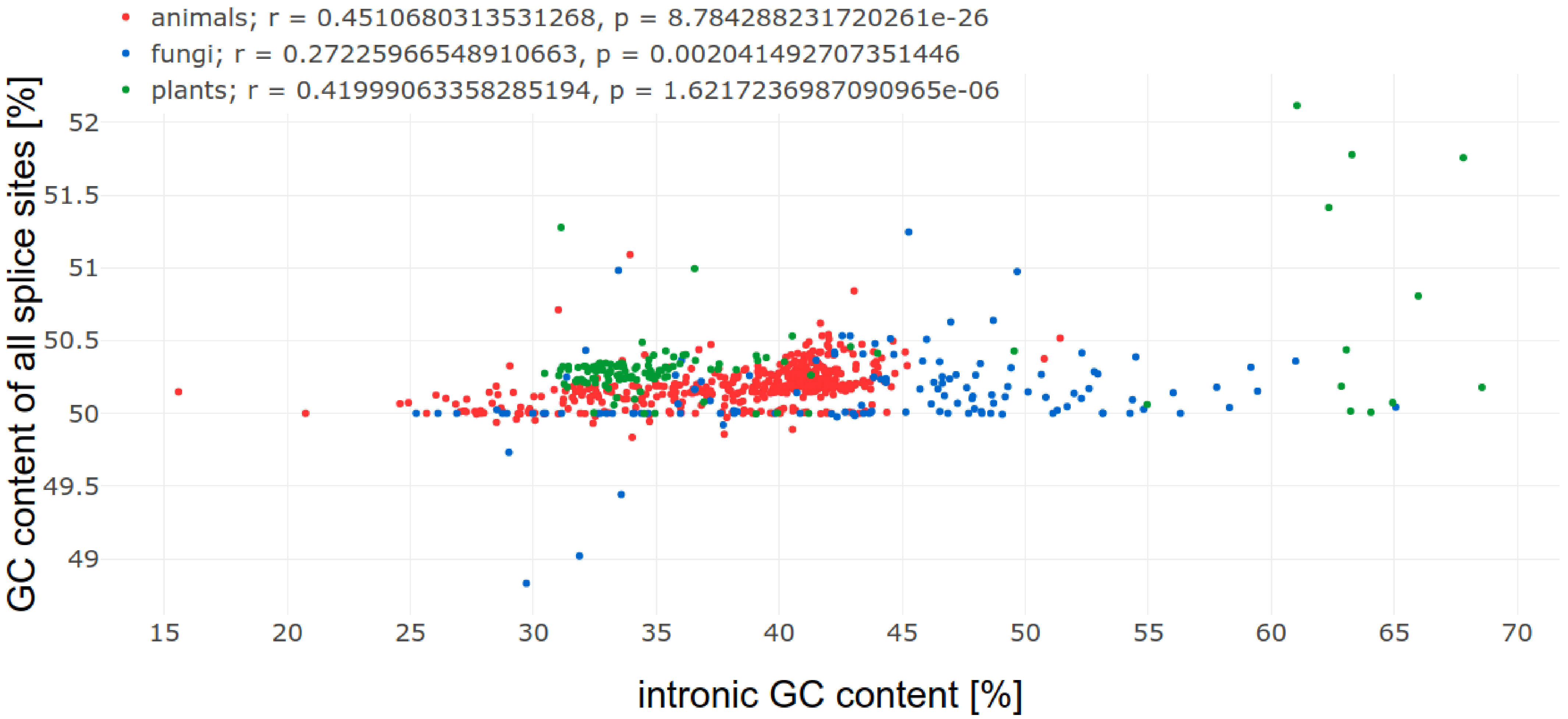

### Supplementary Data S12

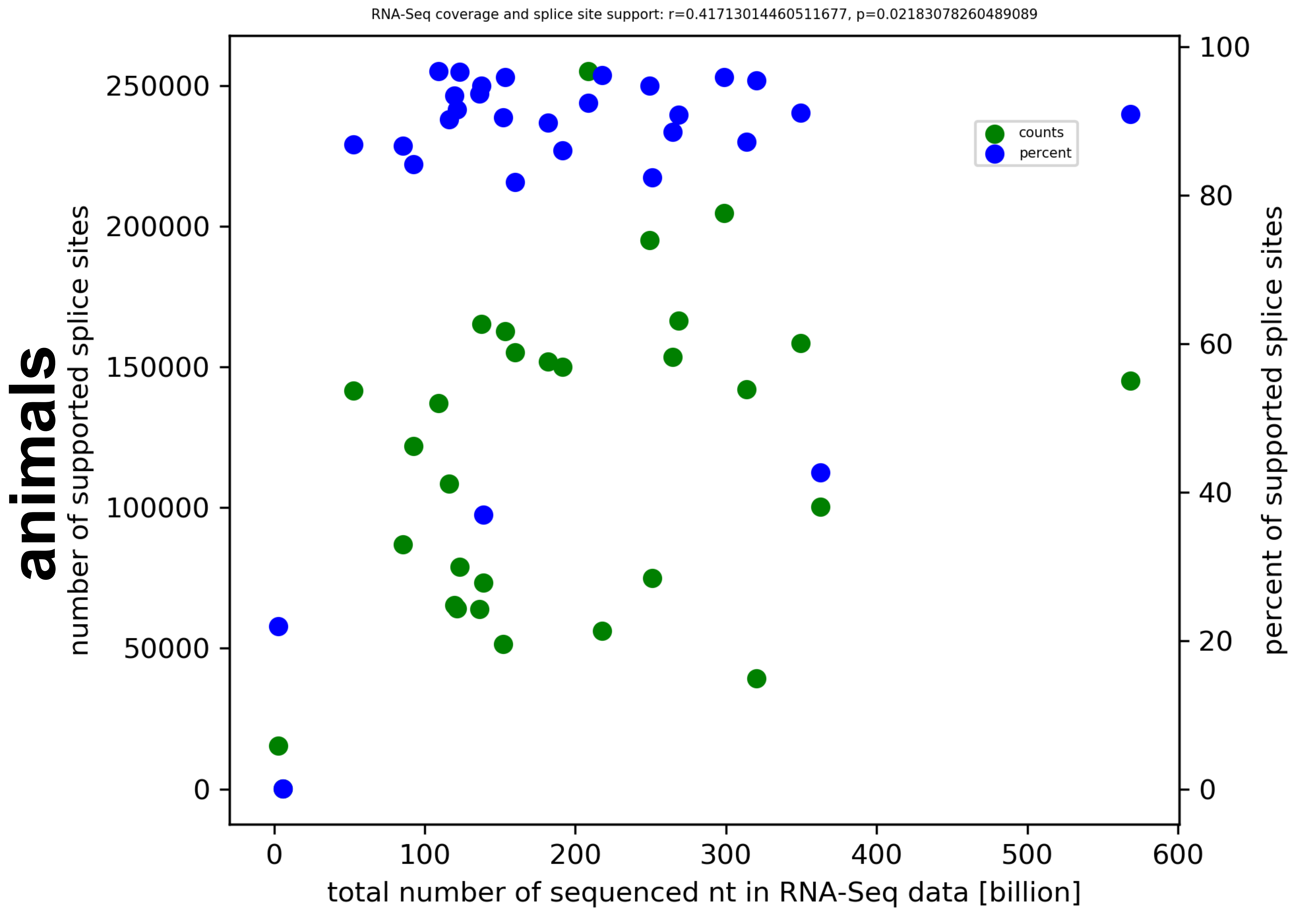

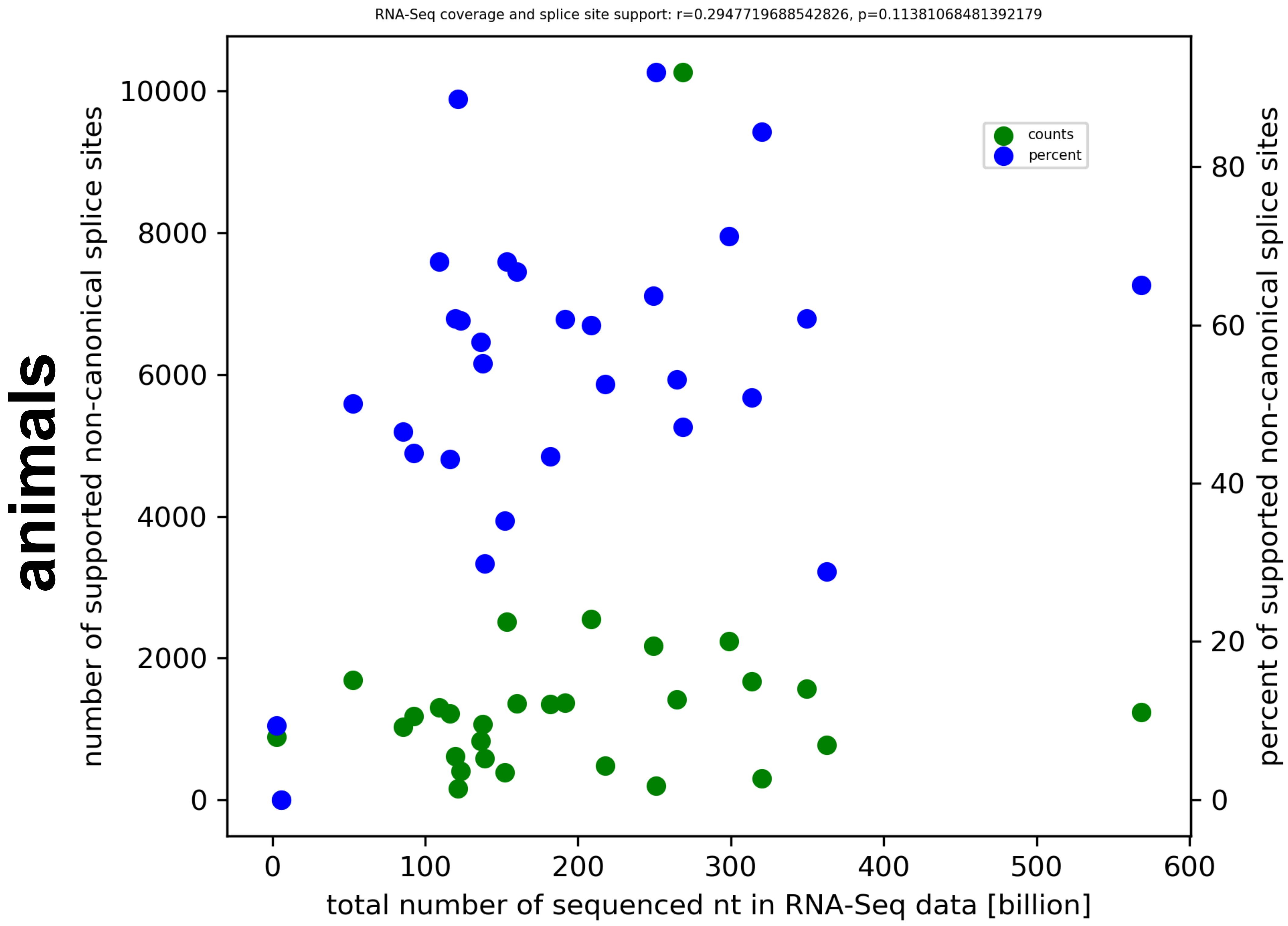

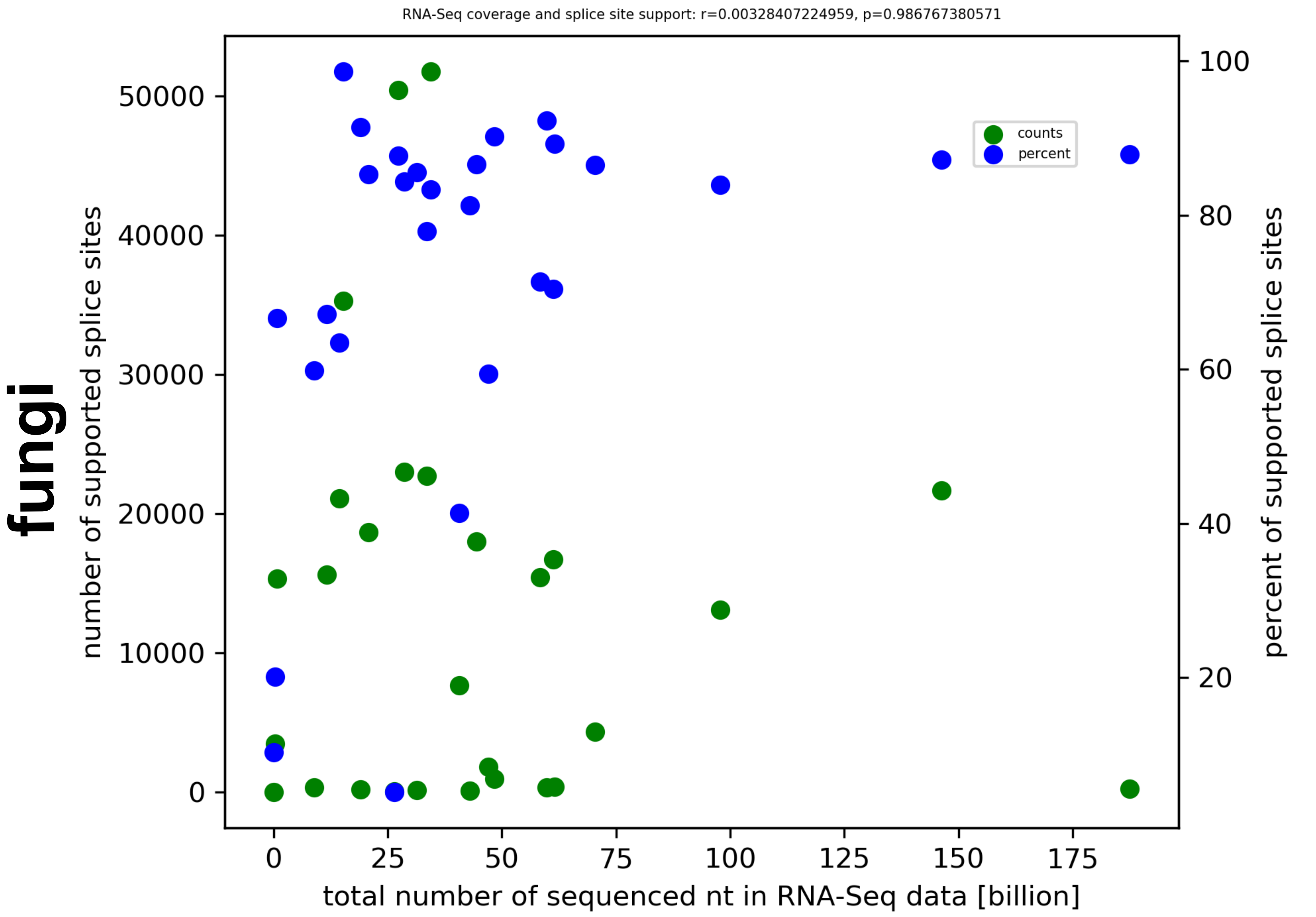

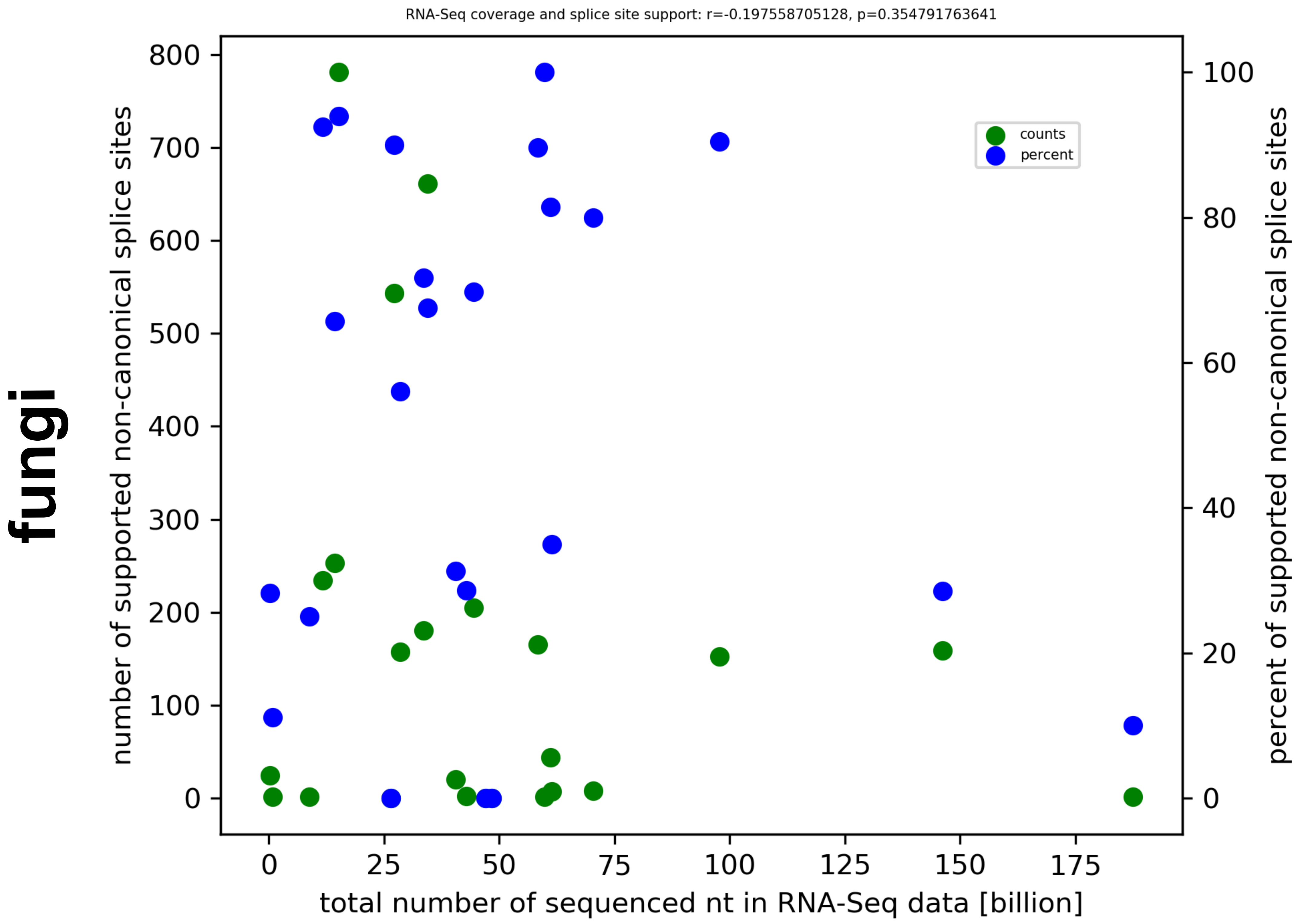

### Supplementary Data S13

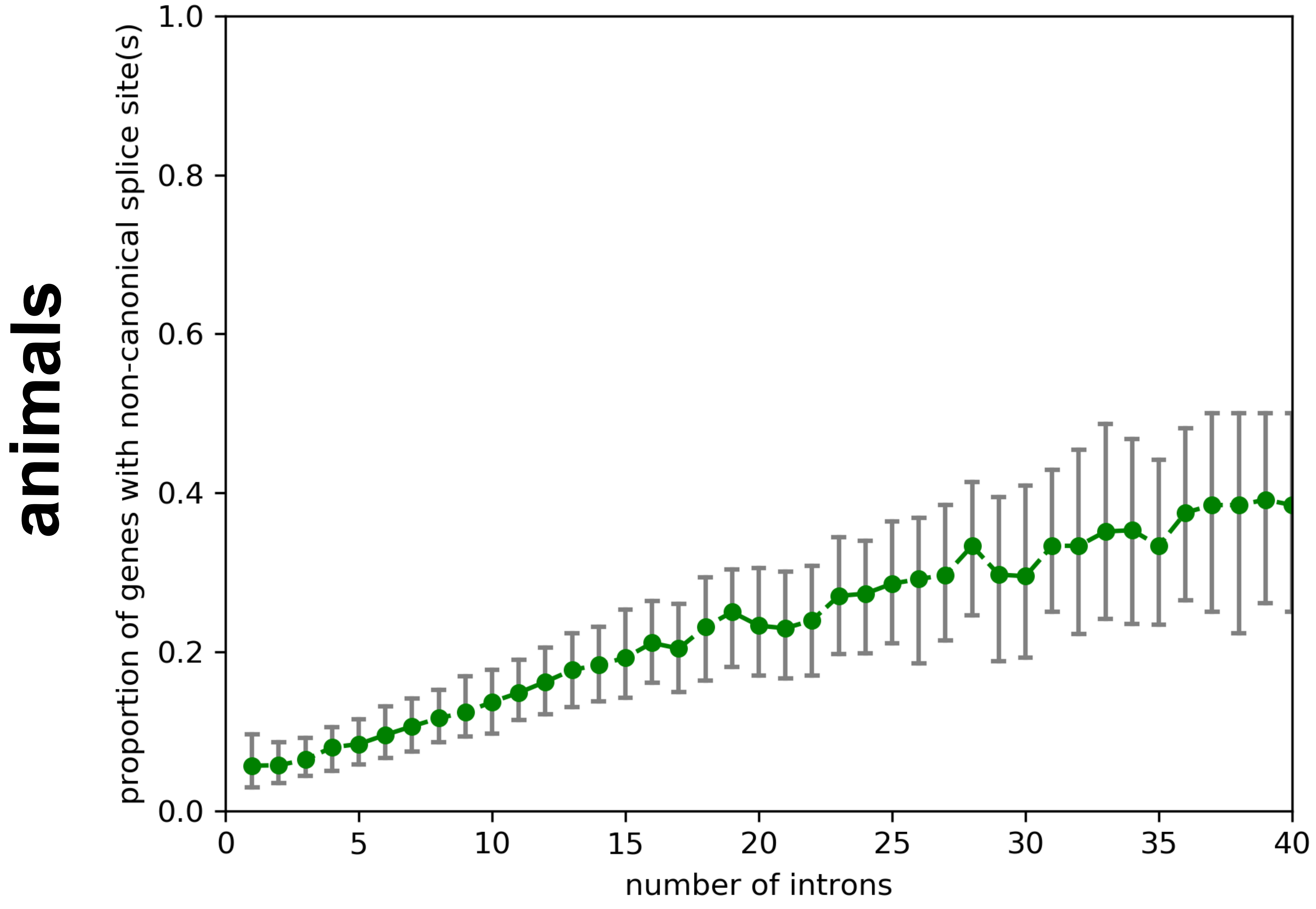

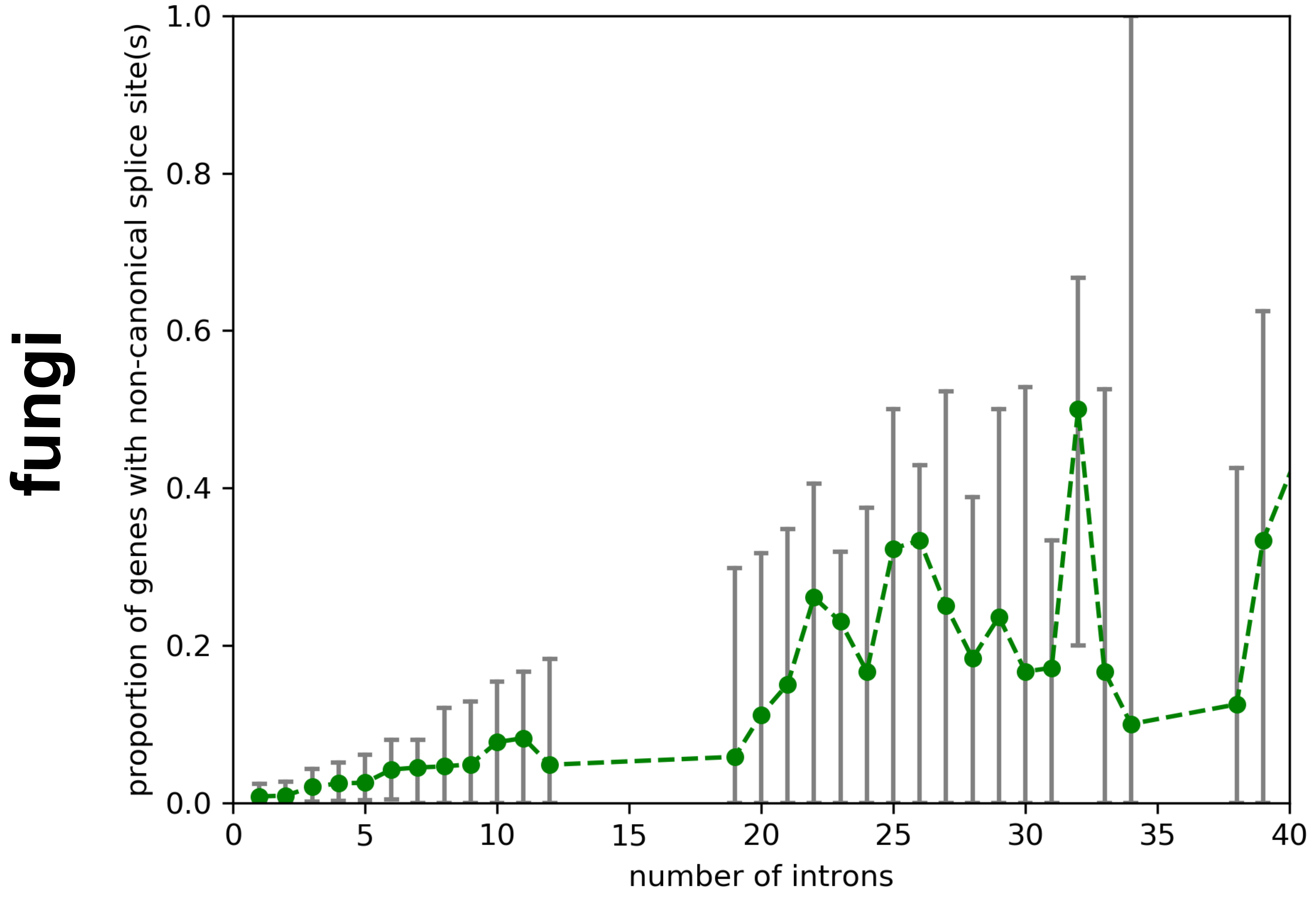

### Supplementary Data S14

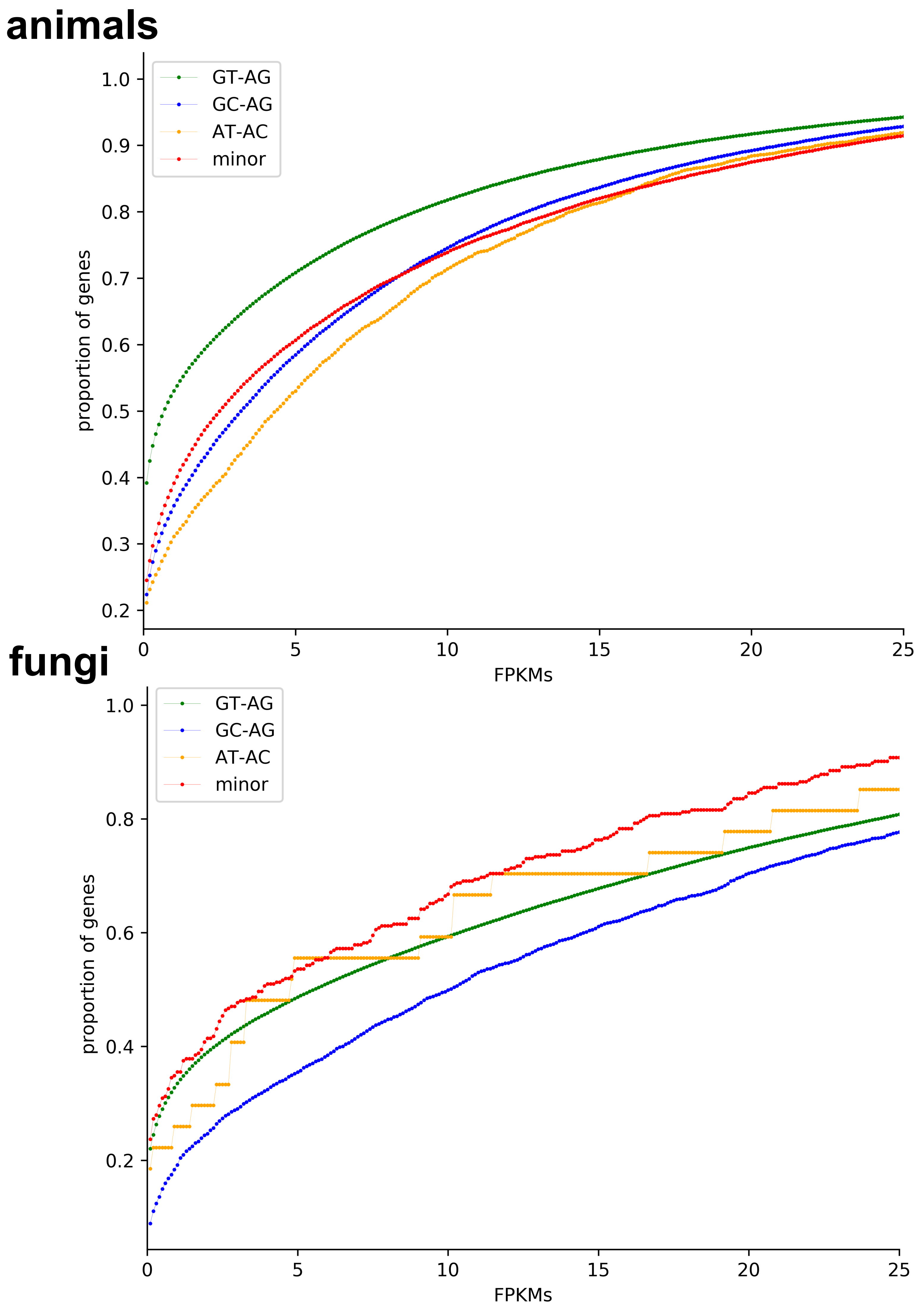

**animals**

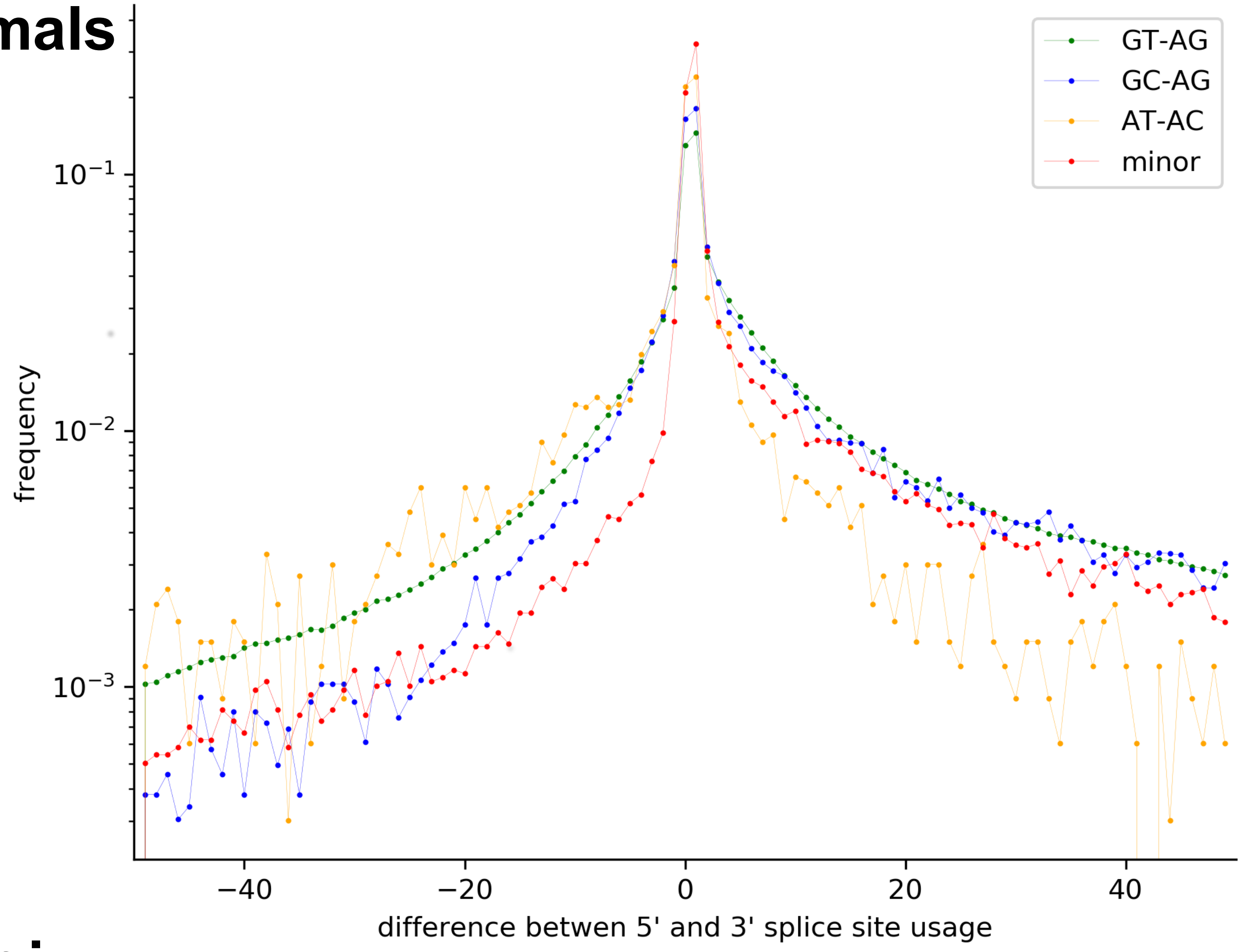


**fungi**

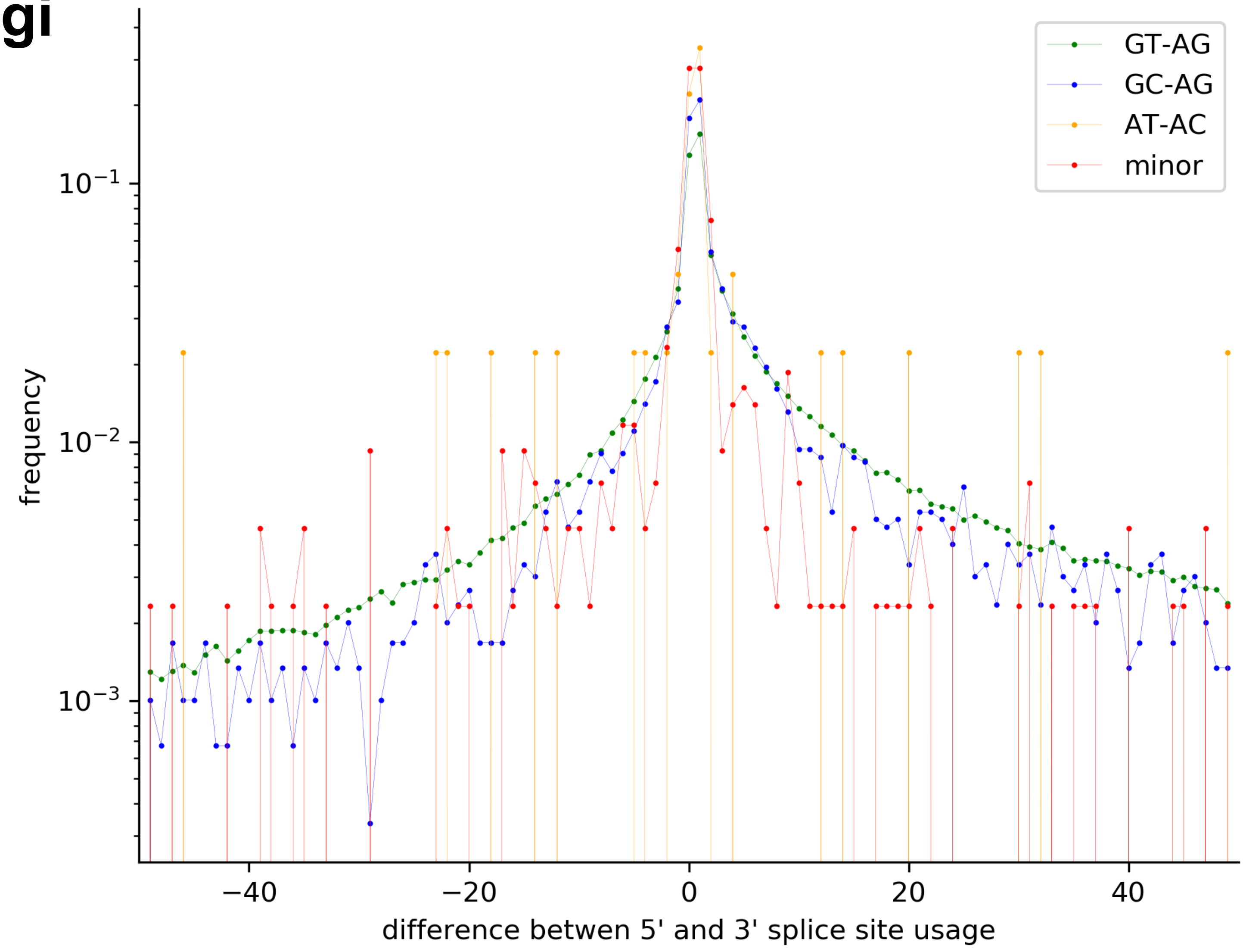
